# Dual transcranial electromagnetic stimulation of the precuneus boosts human long-term memory

**DOI:** 10.1101/2024.10.25.620008

**Authors:** Ilaria Borghi, Lucia Mencarelli, Michele Maiella, Elias P. Casula, Matteo Ferraresi, Francesca Candeo, Elena Savastano, Martina Assogna, Sonia Bonnì, Giacomo Koch

## Abstract

Non-invasive brain stimulation techniques have the potential to improve memory functions. However, the results so far have been relatively modest and time-consuming. Here, we implemented a novel 3-minute combination of personalized repetitive transcranial magnetic stimulation (intermittent theta burst-iTBS) coupled with simultaneous application of gamma transcranial alternating current stimulation (γtACS) over the precuneus, a brain area connected with the hippocampus, to modulate long-term memory in healthy subjects. Only dual electromagnetic stimulation of the precuneus produced an increase in long-term associative memory as compared to iTBS alone and sham conditions in a sample of healthy volunteers. The effects were replicated in another independent sample, in which the increased associative memory was retained for up to one week. Moreover, dual stimulation increased gamma oscillations and precuneus-hippocampus functional connectivity through the white matter tracts linking the precuneus with the temporal lobe. These findings show that dual stimulation may lead neuronal assemblies to a state favorable to enhancing long-term plasticity. Personalized dual electromagnetic stimulation of the precuneus may represent a new powerful approach for enhancing memory functions in several healthy and clinical conditions.

## Introduction

As the world’s population ages, the prevalence of age-related memory deficits is growing. This phenomenon often signals the onset of more severe cognitive decline, which has a strong impact on society. In light of this escalating challenge, there is an urgent need for innovative strategies aimed at enhancing cognitive functions and potentially mitigating cognitive decline.

Non-invasive brain stimulation (NIBS) techniques have been largely used to enhance human cognition in the last two decades (Antal et al., 2022). Encouraging results have been reported in both healthy and pathological conditions including depression, stroke, and Alzheimer’s disease (Blumberger et al., 2018; Koch et al., 2022; Lefaucheur et al., 2020).

Repetitive transcranial magnetic stimulation (rTMS) and transcranial alternating current stimulation (tACS) are two forms of NIBS widely used to enhance memory performance (Grover et al., 2022; Koch et al., 2018; Wang et al., 2014). rTMS, based on the principle of Faraday, induces depolarization of cortical neuronal assemblies and leads to after-effects that have been linked to changes in synaptic plasticity involving mechanisms of long-term potentiation (LTP) (Huang et al., 2017; Jannati et al., 2023). On the other hand, tACS causes rhythmic fluctuations in neuronal membrane potentials, which can bias spike timing, leading to an entrainment of the neural activity (Wischnewski et al., 2023). In particular, the induction of gamma oscillatory activity has been proposed to play an important role in a type of LTP known as spike timing-dependent plasticity, which depends on a precise temporal delay between the firing of a presynaptic and a postsynaptic neuron (Griffiths and Jensen, 2023). Both LTP and gamma oscillations have a strong link with memory processes such as encoding (Bliss and Collingridge, 1993; Griffiths and Jensen, 2023; Rossi et al., 2001), pointing to rTMS and tACS as good candidates for memory enhancement.

However, despite the increasing expectations, their overall effects are relatively modest, even with prolonged stimulation sessions (Grover et al., 2022; Polanía et al., 2018; Wang et al., 2014). In fact, the behavioral and clinical effects are extremely variable (Corp et al., 2020), especially when used to modulate memory functions (Pabst et al., 2022).

Recent studies explored the possibility of combining these two techniques showing that the simultaneous application of tACS in the range of the gamma frequency band (γtACS) with intermittent theta burst (iTBS), an rTMS protocol known to induce LTP (Huang et al., 2005), enhances the after-effects of iTBS alone on the motor and prefrontal cortices (Guerra et al., 2018; Maiella et al., 2022).

Long-term memory formation has been associated with the neural activity of the precuneus (PC). The PC represents one of the main hubs of the default mode network (Cavanna and Trimble, 2006; Cunningham et al., 2017; Jitsuishi and Yamaguchi, 2021) and is involved in the formation of associative, episodic, and autobiographic memories (Bonnì et al., 2015; Brodt et al., 2018; Flanagin et al., 2023). Moreover, it is connected with the temporal lobe and the hippocampus (Cunningham et al., 2017; Dadario and Sughrue, 2023; Tanglay et al., 2022). Hence, the PC represents an ideal target for NIBS to modulate memory functions with profound clinical implications such as in the case of Alzheimer’s disease (Koch et al., 2025, 2024, 2022, 2018).

Here, we hypothesized that γtACS would activate key neural mechanisms underlying memory formation favoring the iTBS-mediated plasticity induction by entraining and synchronizing endogenous gamma oscillations, thereby improving long-term memory. Thus, we reasoned that the dual application of iTBS and γtACS over the PC could be a promising tool for memory enhancement by enhancing plasticity mechanisms.

## Results

### Dual precuneus iTBS+_γ_tACS improves long-term associative memory performance

In the first experiment, subjects (n=20) were asked to perform two tasks measuring long and short-term memory performances immediately after three balanced, randomized, and double-blind stimulation conditions with a cross-over design (iTBS+γtACS, iTBS+sham-tACS, sham-iTBS+sham-tACS). Stimulation parameters were personalized and identified using fMRI and TMS-EEG (see Fig. 1, panels A and B). Long-term memory was assessed by the face-name associative task (FNAT), requiring the memorization of 12 faces with corresponding names and occupations, while short-term memory was assessed by the short-term memory binding test (STMB) (see Fig. 1, panel C).

**Figure 1.**
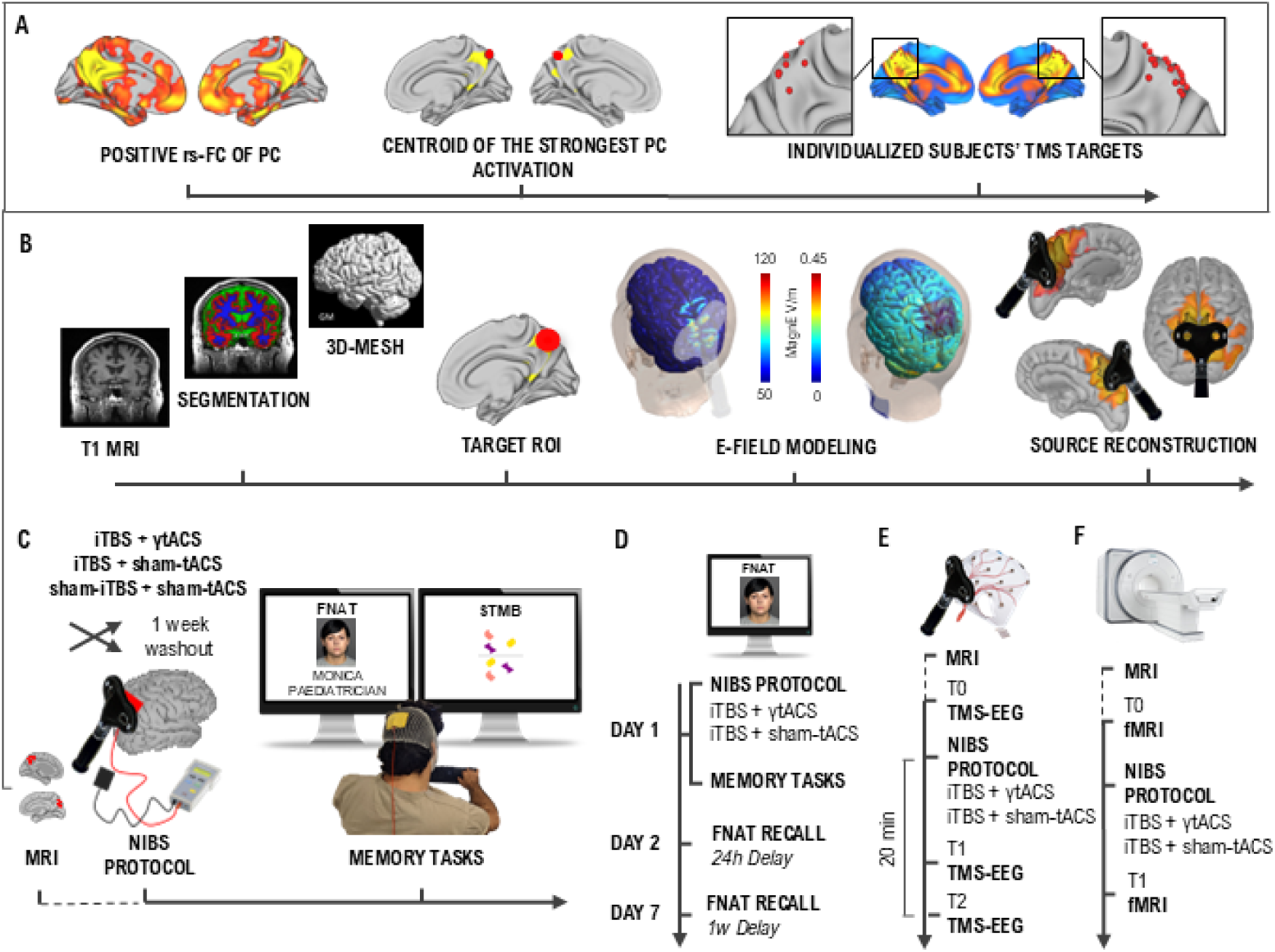
Individual target extraction and experimental design. (**A**) Methodological workflow used to extract the individualized target for the neuromodulation protocol used in experiments 1, 3 and 4. Participants underwent fMRI scanning to individualize the stimulation sites and permit neuronavigation. Target individualization was derived by computing a PC functional connectivity profile for each participant, thus obtaining a map of positively correlated voxels, respectively representing the DMN (Panel A, left). The individual stimulation targets were defined as the centroid of the strongest PC activation being on the top of a cortical gyrus and representing the shortest perpendicular path connecting the stimulating TMS coil on the scalp and the cortex (Panel A, center). PC coordinates for each participant are represented in red on an MNI brain template, showing the overlap with the DMN (Panel A, right). (**B**) Biophysical modeling was computed for each subject acquiring T1w and T2w MRI and using the Simnibs toolbox for the T1w segmentation and the 3D-mesh transformation (Panel B, left). The mean Norm E-field was extracted from a target ROI-sphere (10 mm radius) centered on the individual coordinates of the PC (Panel B, center). The simulated induced electric field is shown for a representative subject produced by iTBS (e-field modeling, left) and tACS (e-field modeling, right). EEG source activity reconstruction induced by the TMS pulse over the precuneus in a representative subject (Panel B, right). Experimental design of experiments 1(**C**), 2 (**D**), 3 (**E**), 4 (**F**). The effect of simultaneous iTBS+γtACS on memory performances was investigated in experiments 1(C) and 2 (D) through two memory tasks: the face-name associative task (FNAT), which required the memorization of 12 faces with corresponding faces and occupations, and the visual short-term memory binding test (STMB), which consisted in a change detection task. In the main experiment 1 (C), subjects were involved in a cross-over design with different experimental sessions of neuromodulation separated by a washout week. Every session corresponded to a different balanced and randomized stimulation condition (i.e., iTBS+γtACS, iTBS+sham-tACS, sham-iTBS+sham-γtACS) immediately followed by the FNAT learning phase and immediate recall, the STMB and the FNAT delayed recall (15-minute delayed) and recognition. In experiment 2 (D) subjects were involved in a cross-over design with two balanced and randomized stimulation conditions (i.e., iTBS+γtACS, iTBS+sham-tACS) separated by a washout week. During the first session (day 1), participants received the neuromodulation protocol and then performed the learning phase and immediate recall FNAT, the STMB and the FNAT delayed recall. In the second session (day 2), participants performed FNAT recall with a 24-hour delay from the neuromodulation protocol, while in the third session (day 7), participants performed FNAT recall and recognition with a 1-week delay. In experiment 3 (E) participants were involved in two randomized and balanced experimental sessions of neuromodulation (i.e., iTBS+γtACS, iTBS+sham-tACS) separated by a washout week. TMS-EEG recordings were performed before (T0), immediately after (T1), and 20 minutes after the neuromodulation (T2). In experiment 4 (D), after the first MRI scanning used for neuronavigation, participants were involved in two randomized and balanced experimental sessions of neuromodulation (i.e., iTBS+γtACS, iTBS+sham-tACS) separated by a washout week. The fMRI scanning was performed before (T0) and immediately after (T1) the neuromodulation protocol. Photographs reported represent the author Michele Maiella performing the task and an example of the face item used in the task taken from the FACES database (Ebner et al., 2010).

The different stimulation protocols were all well-tolerated. No one reported significant side effects connected with the neuromodulation protocol applications. We found that dual iTBS+γtACS improved long-term associative memory as compared to the other conditions. Dual iTBS+γtACS increased the performance in recalling the association between face, name and occupation (FNAT accuracy), both for the immediate (F_2,38_=7.18; p=0.002; η²_p_=0.274) and the delayed (F_2,38_=5.86; p=0.006; η²_p_=0.236) recall performances (Fig. 2, panel A). The sensitivity analysis showed a minimal detectable effect size of η²_p_=0.215 with 20 participants. Post-hoc analysis showed an advantage for dual iTBS+γtACS as compared to iTBS+sham-tACS condition for the immediate (40.0±21.6% vs. 25.0±17.9%, p=0.002 Bonferroni corrected, corresponding to a relative improvement of 63.79%) and for the delayed recall (34.2±20.4% vs. 24.2±19.8%, p<0.05 Bonferroni corrected, corresponding to a relative improvement of 34.16%), as well as compared to the sham-iTBS+sham-tACS condition for the delayed recall (34.2±20.4% vs. 26.3±17.6% p=0.037 Bonferroni corrected, corresponding to a relative improvement of 34.04%) (Fig. 2, panel A).

**Figure 2.**
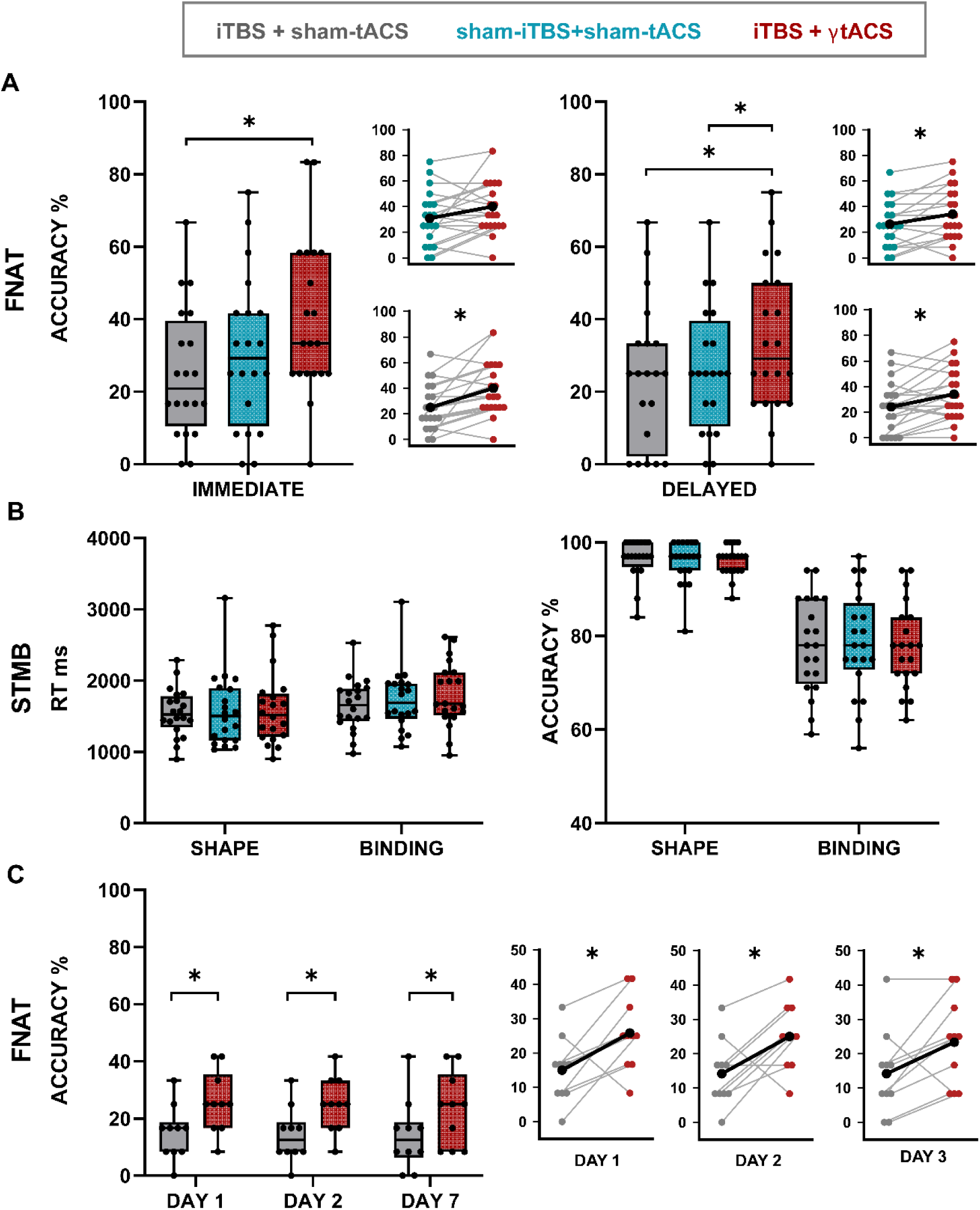
Memory performance outcome. In the Box and Whiskers plots, boxes delimit the lower (Q1) and the upper (Q3) quartile, the whiskers extend to the smallest and largest values, dots represent individual values. In the line graphs, colored dots show individual participant data, lines show the connection between the performances of each participant, black dots and lines correspond to the mean performance values. Grey corresponds to iTBS+sham-tACS, blue to sham-iTBS+sham-tACS and red to iTBS+γtACS. * = p<0.05 (**A**) Face-name associative task (FNAT) accuracy in immediate (right) and delayed (left) trials resulting from experiment 1. N=20. (**B**) Short-term memory binding task RTs (left) and accuracy (right) resulting from experiment 1. N=20. (**C**) FNAT’s long-lasting effect resulted from experiment 2. The results are shown over time (day 1, day 2, day 7). N=10.

The in-depth analysis of the FNAT accuracy investigating the specific contribution of face-name and face-occupation recall revealed that dual iTBS+γtACS increased the performance in the association between face and name (FNAT NAME) delayed recall (F_2,38_ =3.46; p =0.042; η²_p_ =0.154; iTBS+γtACS vs. sham-iTBS+sham-tACS: 42.9±21.5 % vs. 33.8±19 %; p=0.048 Bonferroni corrected) (Fig. S4, supplementary information). iTBS+sham-tACS did not result in any improvement as compared to sham-iTBS+sham-tACS. No significant effects were found over the recall of the association between face and occupation (FNAT OCCUPATION) (p<0.05). Moreover, we found that the effects of dual iTBS+γtACS stimulation were selective for long-term but not short-term memory, since no effects were found on STMB accuracy and RTs (all ps>0.05) (Fig. 2, panel B). Table S1 (supplementary information) shows Experiment 1 statistical details. Biophysical modeling calculations showed that the iTBS induced a mean e-field of 29.55±5.17 V/m, while γtACS led to a much smaller mean e-field of 0.1±0.55 V/m.

### Enhanced memory performance is still evident after one week

To confirm the above-described effects of dual iTBS+γtACS on an independent sample (n=10) and to investigate the time course of memory retention we performed a second experiment. In this experiment, subjects were asked to execute the delayed FNAT cued recall 24 hours and a 1-week after the neuromodulatory protocols. As for experiment 1, we confirmed that dual iTBS+γtACS improved immediate recall at TOTAL FNAT as compared to iTBS+sham-tACS (F_1,9_=7.31; p=0.024; η²_p_=0.448; 26.7±10.2% vs. 17.5±8.3%; p=0.024 corresponding to a relative improvement of 82.50%) (see Table S2 in supplementary information for statistical details). The sensitivity analysis showed a minimal detectable effect size of η²_p_=0.388 with 10 participants. More importantly, we found that dual iTBS+γtACS exerted a long-lasting effect on long-term associative memory (F_1,9_=8.433; p=0.017; η²_p_=0.484) (Fig. 2, panel C). The sensitivity analysis showed a minimal detectable effect size of η²_p_=0.125 with 10 participants. We observed that the incremental effect on TOTAL FNAT performance was still evident at 24 hours and one week after the neuromodulation (day 1: 25.8±10.7 % vs. 15±9.5 %, t_9_ =-2.75, p=0.011 corresponding to a relative improvement of 78.33%; day 2: 25±9.6 % vs. 14.2±9.7 %, t_9_=-2.89, p=0.009 corresponding to a relative improvement of 88.33%; day 7: 23.3±12.9 % vs. 14.2±12.5 %, t_9_=-2.40, p=0.020 corresponding to a relative improvement of 80%) (see Table S3 in supplementary information). No significant effects were found for STMB accuracy and RTs (all ps>0.05) as for experiment 1 (see table S2 in supplementary information for statistical details).

### Precuneus iTBS+_γ_tACS increases gamma oscillatory activity

We then aimed to investigate the effects of the neuromodulation protocol on cortical oscillations (TRSP) and excitability (TEPs), through TMS-EEG recordings (n=14 subjects, 6 of whom also participated in experiment 1). We found that cortical oscillations in the gamma band increased after dual iTBS+γtACS as compared to baseline TMS-EEG recordings (stimulation F_1,13_=5.073; p=0.042; η²_p_=0.281) (Fig. 3, panel A). Specifically, we observed an increase in the gamma-TRSP after dual iTBS+γtACS as compared to iTBS+sham-tACS in both time points (ΔT1: 0.045±0.09 vs. -0.013±0.06; t_13_=1.96; p=0.036; ΔT2: 0.026±0.04 vs. -0.013±0.05; t_13_=1.92; p=0.039) (Fig. 3, panel B). This effect was specific to PC; indeed, no significant effect was found when testing TMS-EEG over the l-PPC (p>0.05). This result was reflected by increased cortical excitability evident after dual iTBS+γtACS but not after iTBS+sham-tACS condition (Fig. 3, panel C). Specifically, we observed a significant TEP amplitude increase over a cluster of parietal and occipital electrodes comparing T0 vs T1 (mean t-value_13_=-3.29; all ps<0.05) and over POz (t-value_13_=-4.74; p<0.05) when comparing T0 vs. T2. No significant differences were found comparing T0 vs. T1 and T2 in the iTBS+sham-tACS condition (ps>0.05).

**Figure 3.**
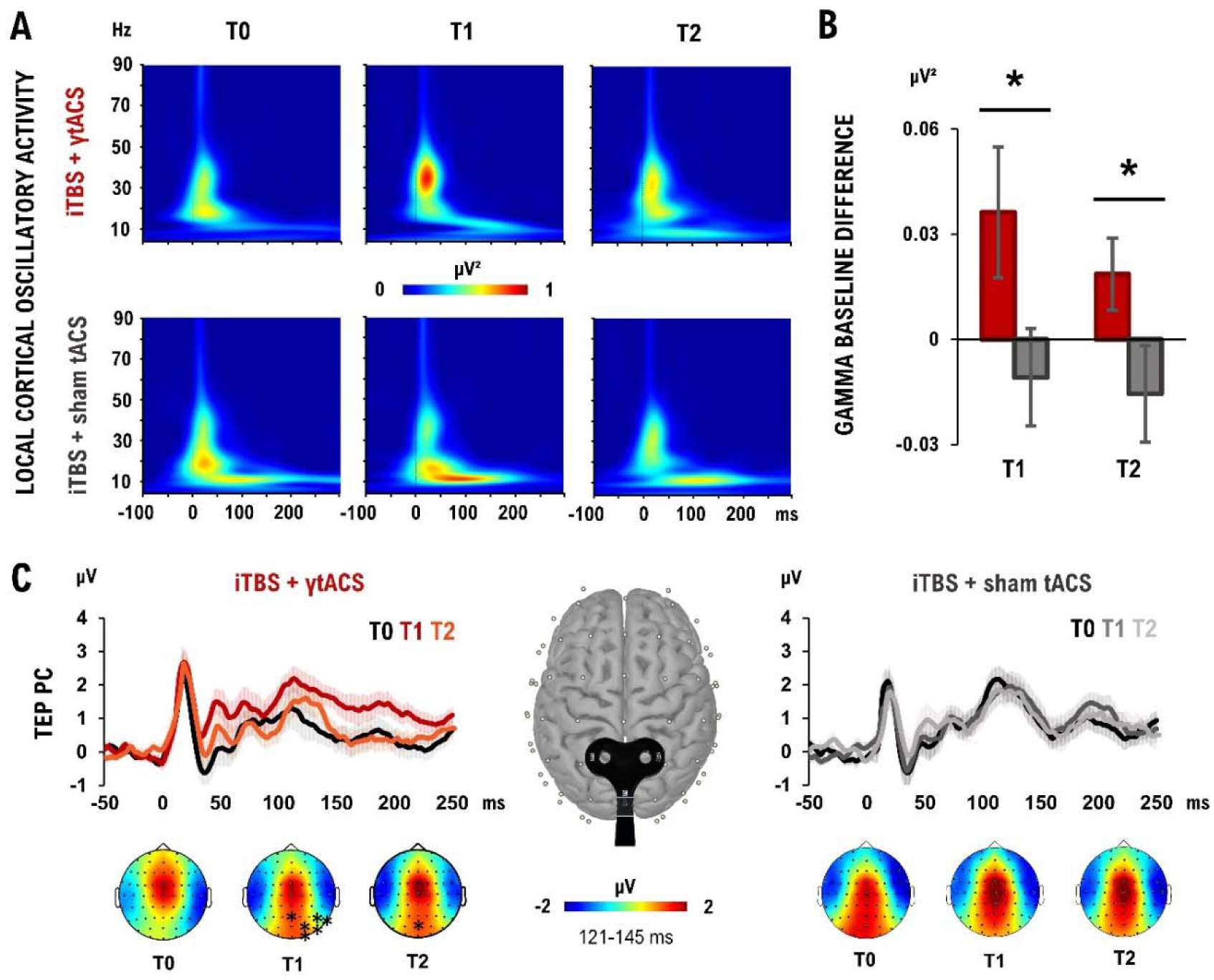
Experiment 3 neurophysiological outcome. (A) Precuneus (PC) oscillatory activity elicited by iTBS+γtACS (up) and iTBS+sham-tACS (down) when testing PC over the three time points (T0, T1, T2 from left to right). (B) Gamma oscillation changes from baseline after iTBS+γtACS (red) and iTBS+sham-tACS (grey). (C) TMS-evoked potential (TEP) produced over the PC when performing TMS-EEG over PC in the two stimulation conditions: iTBS+γtACS (up-left) and iTBS+sham-tACS (up-right) over the three time points (T0, T1, T2). (Down) Topographies and statistical differences in TEPs amplitude after the different stimulation conditions (iTBS+γtACS, left; iTBS+sham-tACS, right) over the three time points (T0, T1, T2, from left to right). N=14; * = p<0.05; bars depict standard error.

### iTBS+_γ_tACS increases precuneus connectivity with the hippocampus

Finally, we investigated the effects of the dual iTBS+γtACS protocols on MRI-based resting state functional connectivity (n=16 subjects, 7 of whom participated in experiment 1). ROI-to-ROI analysis revealed a significant difference specifically for the connectivity between the PC and the hippocampi (F_3,13_= 5.78, p-FDR= 0.029). Post-hoc analysis showed increased connectivity after the stimulation for the dual iTBS+γtACS (t_15_ =3.67, p-FDR=0.0023; Fig. 4, panel A), but not for the iTBS+sham-tACS. No significant differences were detected among the conditions before stimulation (all p>0.05). The significant nodes that emerged from the ROI-to-ROI analysis (i.e., PC and hippocampi) were considered for the seed-to-voxel analyses, revealing an increased rs-FC between the PC and the orbitofrontal cortex after dual iTBS+γtACS as compared to baseline (OFC; MNI coordinates: x=20, y=16, z=-4; |T15| > 3.20, k=1051; p=0.0003 (Fig. 4, panel B center). Moreover, stronger connectivity with the PC (MNI coordinates: x=30, y=-70, z=46; |T15| > 3.20, k=1499; p=0.000167) and with the posterior cingulate cortex emerged when considering the right hippocampus as seed (MNI coordinates: x=-14, y=-50, z=12; |T15| > 3.20, k=1295; p=0.0001649) (Fig. 4, panel B right). Finally, similar results occurred considering as seed the left hippocampus, with an improved rs-FC with the PC (MNI coordinates: x=30, y=-68, z=44; |T15| > 3.20, k=1525; p=0.000163) and with the posterior cingulate cortex (MNI coordinates: x=20, y=-50, z=12; |T15| > 3.20, k=509; p=0.022) (Fig. 4, panel B left). We also reconstructed the white matter fibers connecting the PC with the temporal lobe in each subject, forming the bilateral Middle Longitudinal Fasciculus (MdLF), using DTI tractography. We found that the structure of the MdLF measured through fractional anisotropy (FA) was related to the increased connectivity between PC and the hippocampi after dual iTBS+γtACS. A significant positive correlation emerged between the FA of the MdLF and the changes in connectivity between PC and hippocampus in the iTBS+γtACS (r=0.73, p=0.001), but not in the iTBS+sham-tACS (r= 0.14, p=0.634) (Fig. 4, panel C).

**Figure 4.**
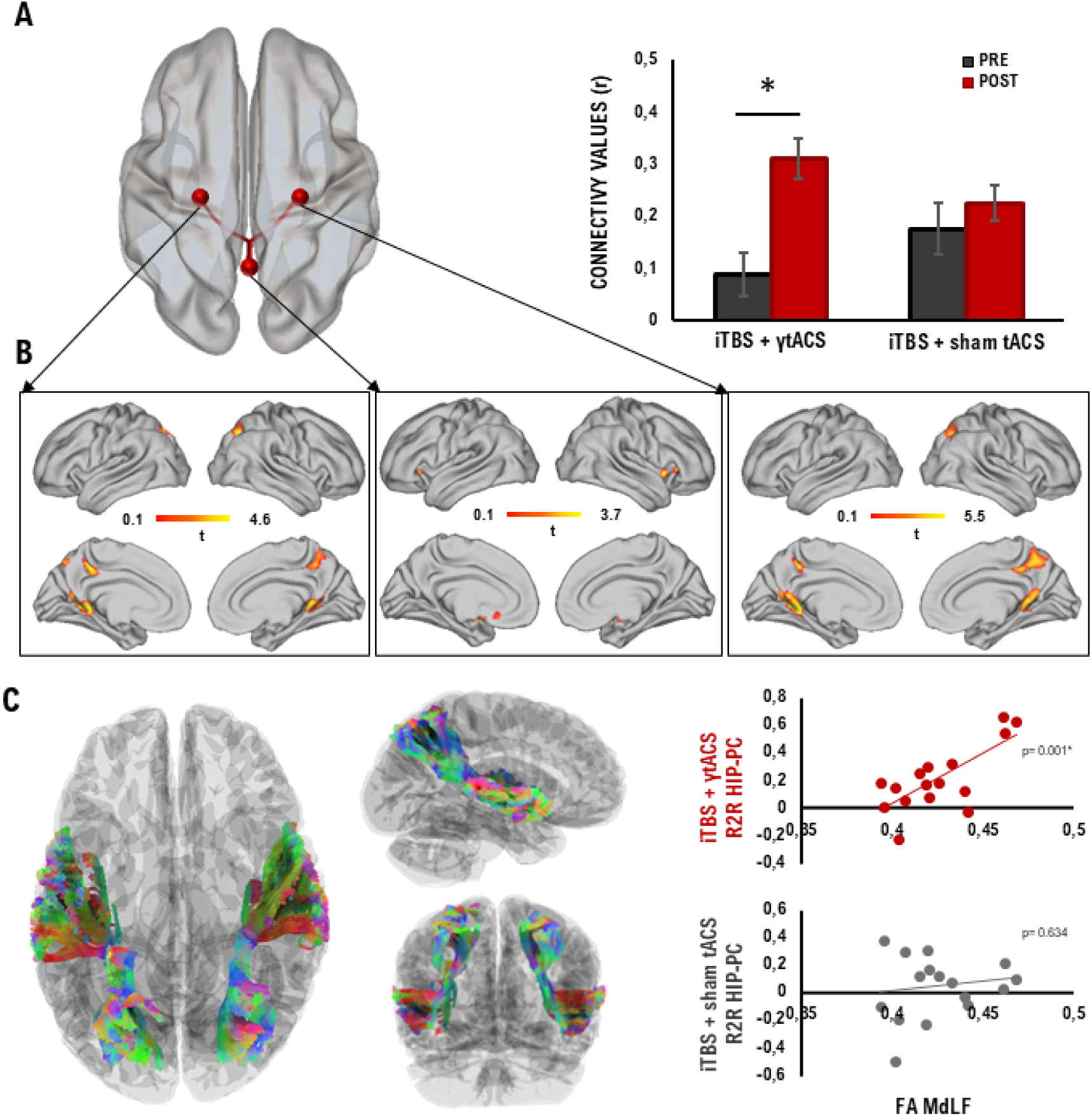
**rsFC changes after iTBS+**_γ_**tACS and correlation with DTI.** (A) The standard MNI brain (left) and the bar plot (right) display the positive correlation between the PC and the bilateral HIP after the iTBS+γtACS resulted from the ROI-to-ROI analysis. (B) Seed-to-voxel analysis results from each significant ROI (i.e. left HIP, left; PC, center; right HIP, right) are overlaid to a standard MNI brain. (C) MdLF extracted (left); positive correlation between MdLF integrity and functional connectivity changes between the PC and bilateral HIP after iTBS+γtACS (up right); the absence of correlation in the iTBS+sham-tACS condition and functional connectivity (down right). N=16; * = p<0.05; bars depict standard error.

## Discussion

Here, we demonstrated that the combination of iTBS and γtACS applied over the personalized PC can enhance long-term associative memory formation. Dual stimulation increased the ability to recall the association between faces, names, and occupations as compared to the iTBS alone or sham conditions. Moreover, the increased memory trace was long-lasting, being evident one week after the stimulation. On the other hand, iTBS alone did not result in any memory improvement, in agreement with previous studies showing contradictory effects of this protocol on neuroenhancement (Hamada et al., 2013; Ziemann and Siebner, 2015). Hence, we show that the combination of neuroplasticity induction promoted by iTBS and gamma oscillations entrainment produced by γtACS, may be the key to memory boosting. We believe that the novel results reported here may have several implications for treating memory impairments, gamma dysregulation, and network dysfunction in a large variety of conditions.

Our results seem to identify the PC as having a role in mediating long-term memory formation. Given the existing evidence on TMS propagation and the computation of the biophysical model with the E-field, we can assume that the individually identified PC was involved in the observed effects (Ridding and Rothwell, 2007). Moreover, we observed specific cortical changes in the posteromedial parietal areas, as evidenced by the whole-brain analysis conducted on TMS-EEG data and the absence of effect on the lateral posterior parietal cortex used as a control condition. The PC has been associated with memory engram formation and retrieval (Brodt et al., 2018; Flanagin et al., 2023; Hebscher et al., 2019). Moreover, it is a prime candidate in processing relations among entities, including semantic concepts, making its activity fundamental for associative memory formation (Fernandino et al., 2022; Rentz et al., 2011; Summerfield et al., 2020). These notions found support in our current findings, showing that neuromodulation of the PC resulted in an improvement of long-term associative memory.

Visual short-term associative memory, measured by STBM performance, was not modulated by any experimental condition. Even if we cannot exclude the possibility that the stimulation could have influenced short-term verbal associative memory, we expected this result since short-term associative memory is known to rely on a distinct frontoparietal network while FNAT, used to investigate long-term associative memory, has already been associated with the neural activity of the PC and the hippocampus (Parra et al., 2014; Rentz et al., 2011). Moreover, we observed a main effect on the ability to concomitantly recall face, name and occupation, while for single association, we observed minimal (face-name) or no effect (face-occupation). Forming face-name associations is widely acknowledged to be particularly difficult due to the lack of contextual properties to formulate an associative link (Werheid and Clare, 2007). In contrast, forming an association between a face and other biographical information, such as occupations or hobbies, is easier (McWeeny et al., 1987). We can hypothesize that retaining multiple associations (face-name-occupation) together is even more effortful and cognitively demanding. For these reasons, the stimulation could have promoted effects in a gradient way related to the difficulty of the task. Apart from improving long-term memory, dual iTBS+γtACS promoted a long-term increase in gamma oscillatory activity, as measured by TMS-EEG. Cortical gamma-band oscillations are thought to be primarily generated by parvalbumin-positive GABA-ergic interneurons interacting with excitatory pyramidal cells (Buzsáki and Wang, 2012; Cobb et al., 1995). The repeated activation of these circuits through gamma oscillations fosters LTP formation, which is the neurobiological basis of long-term memory (Bikbaev and Manahan-Vaughan, 2008; Debanne and Inglebert, 2023; Headley and Weinberger, 2011). This finding is consistent with the idea that increased gamma oscillatory activity may facilitate cortical plasticity, neural communication, and mechanisms of memory encoding and retrieval, thereby ensuring the successful formation of an episodic memory (Griffiths and Jensen, 2023). We hypothesize that γtACS, by entraining and synchronizing ongoing oscillatory gamma activity, may have facilitated the formation of LTP induced by iTBS.

Furthermore, we found that dual iTBS+γtACS enhanced the connections between the PC and the hippocampus as measured by fMRI connectivity analysis. Effective neural communication driven by gamma oscillations ensures that, during memory formation, incoming information processed in the neocortex activates the relevant cell assemblies in the hippocampus to ensure associative binding (Griffiths and Jensen, 2023). Gamma-band oscillatory activity drives network connectivity, allowing for information processing and binding through synchronization between distant cortical areas (Fries, 2005; Jensen et al., 2007; Miltner et al., 1999; Osipova et al., 2006). Hence, we argue that PC gamma oscillations could have bound relevant stimuli for perceptual representation, while synchronization between cortical and hippocampal neurons could have allowed the representation to be encoded into the hippocampus (Nyhus and Curran, 2010).

Finally, our findings seem to suggest that higher functional connectivity was supported by higher white matter integrity (FA) of MdLF, the main white matter tract connecting the PC with the temporal lobe. Altogether, these results help to trace the possible anatomo-functional network engaged by PC stimulation, providing further support for the successful involvement of long-range pathways connecting the PC with the temporal lobe underlying memory formation.

A strength of the current work is the personalization of stimulation parameters that was achieved by targeting the PC spot more connected with the temporal lobe and by selecting the intensity of stimulation for iTBS using TMS-EEG. Hence, we argue that personalization of NIBS protocols is a key factor in reducing the inter-individual variability that currently limits the use of common protocols such as TBS and in increasing the overall beneficial effects on cognitive functions.

Why should the dual application of γtACS+iTBS lead to such a strong boosting effect as compared to iTBS alone? It is well established that tACS alone does not induce relevant after-effects, especially when applied for a short period of 3 minutes, such as in the current case (Guerra et al., 2019, 2018). However, during stimulation, tACS alters membrane potential towards depolarization or hyperpolarization in an oscillatory fashion, manipulating the excitability of neurons that become aligned with the introduced electric field, mostly pyramidal cells in layer V (Fröhlich and McCormick, 2010; Guerra et al., 2018; Reato et al., 2013; Schutter and Hortensius, 2011). Notably, these neurons are characterized by intrinsic resonance and neuroplastic activity. In particular, synchronous neural activation in the γ range is considered crucial for the induction of neuroplasticity and memory formation (Buzsáki and Wang, 2012). Gamma oscillations can synchronize the firing of multiple presynaptic neurons so that they exert a stronger depolarizing effect on the target postsynaptic neuron than if they were to fire in isolation (Sjöström et al., 2001). Synchronizing multiple inputs to a postsynaptic neuron enhances the likelihood of LTP. Hence, gamma oscillations are perhaps ideal because they provide a comparatively short window of excitability that ensures all neurons fire in near-perfect unison (Jensen et al., 2007). This provides a long-sought link between gamma oscillations, LTP, and the formation of new memories. For these reasons, γtACS may have provided the ideal neural substrate for the iTBS to exert its plasticity-inducing mechanisms. This might have occurred by driving neuronal assemblies in a state devoted to inducing long-term changes in cortical networks involved in memory formation. In humans, iTBS has been inspired by protocols used in animal models to elicit LTP in the hippocampus (Huang et al., 2005).

On top of iTBS, γtACS may have synchronized PC neuronal elements, boosting gamma oscillatory activity during the induction of LTP driven by iTBS. Notably, this interaction is rather specific, since previous studies showed that the application of γtACS alone does not induce any after-effects, while only gamma, but not theta or beta tACS, combined with iTBS induces long-lasting changes in cortical excitability (Guerra et al., 2018; Maiella et al., 2022). Moreover, our biophysical modeling calculations showed that γtACS induced a marginal augmentation of the e-field (∼0.15 mV), which is below the amount of induced current needed to induce an effect in superficial cortical neurons (Opitz et al., 2016). Hence, tACS alone cannot account for the observed increase in memory as well as for enhancing cortical activity.

This study has some limitations. Although we studied TMS and tACS propagation through the E-field modeling and observed an increase in the precuneus gamma oscillatory activity, excitability and connectivity with the hippocampi, we cannot exclude that our results might reflect the consequences of stimulating more superficial parietal regions other than the precuneus nor report direct evidence of microscopic changes in the brain after the stimulation. Invasive neurophysiological recordings in humans for this type of study are not feasible due to ethical constraints. Studies on cadavers or rodents would not fully resolve our question due to significant differences between them (i.e., rodents do not have an anatomical correspondence, while cadavers have alterations in electrical conductivity occurring in postmortem tissue). However, further exploration of this aspect in future studies would help in the understanding of γtACS+iTBS effects.

We did not assess the effects of γtACS alone. This decision was based on the findings of Guerra et al. (Guerra et al., 2018), who investigated the same protocol and reported no aftereffects. Given the substantial burden of the experimental design on patients and our primary goal of demonstrating an enhancement of effects compared to the standalone iTBS protocol, we decided to leave out this condition. While examining the effects of γtACS alone could help isolate its specific contribution to this target and memory function, extensive research has shown that achieving a cognitive enhancement aftereffect with tACS alone typically requires around 20–25 minutes of stimulation (Grover et al., 2023). We did not study memory functions, gamma oscillations, and synchronization between the PC and hippocampus in the same session due to technical limitations with the current techniques (i.e., fMRI, TMS-EEG). Consequently, these findings do not allow precise inferences regarding the specific mechanisms by which dual iTBS and γtACS of the precuneus modulate learning and memory. Methodological restraints related to the high-frequency protocols employed in this study did not allow for closed-loop implementation. Moreover, we did not test tACS applied at different frequency bands combined with iTBS since our working hypothesis was based on the combination of gamma activity and plasticity induction. Future studies should further investigate the effects of stimulation on distinct memory processes. In particular, stimulation could be applied before retrieval (Rossi et al., 2001) to better elucidate its specific contribution to the observed enhancements in memory performance. Additionally, it would be worth examining whether repeated stimulation, administered both before encoding and before retrieval, could produce a boosting effect. This is especially relevant in light of findings showing that matching the brain state between retrieval and encoding can significantly enhance memory performance (Javadi et al., 2017).

### Our findings may have profound social and clinical implications

These results may assume relevance in the context of memory impairment, which is an increasing and challenging aspect of the elderly population. The current findings may be relevant in the case of Alzheimer’s disease, where the progressive memory deficit is accompanied by early involvement of DMN with an accumulation of beta-amyloid plaques, neurofibrillary tangles, atrophy, and functional connectivity primarily targeting the PC (Billette et al., 2022; Buckner et al., 2005; Chen et al., 2017; Raichle et al., 2001). In this perspective, the DMN has been recently identified as a new potential therapeutic target for neuromodulation in AD (Koch et al., 2025, 2024, 2022, 2018). Moreover, by linking together items and concepts, long-term associative memory is essential for the construction of declarative memory and, broadly for learning and everyday functioning. Hence, dual magnetic and electrical transcranial stimulation could be promising in several social and clinical contexts, such as learning deficits, autism spectrum, and attention deficit hyperactivity disorders (Mathalon and Sohal, 2015).

## Materials and Methods

### Experimental design

We conducted four experiments aimed at understanding the effects of dual iTBS+γtACS on associative memory performance, cortical oscillatory activity and reactivity, and functional connectivity.

*Experiment 1.* We investigated the effect of dual iTBS+γtACS on memory performance through two memory tasks: the face-name associative task (FNAT) and the visual short-term memory binding test (STMB). Participants first underwent MRI scanning to individualize the stimulation sites and permit neuronavigation (Fig. 1, panel A), then they were involved in a double-blind cross-over design with three experimental sessions of neuromodulation separated by a washout week (Fig. 1, panel C). Every session corresponded to a different balanced and randomized stimulation condition (i.e., iTBS+γtACS, iTBS+sham-tACS, sham-iTBS+sham-γtACS) immediately followed by the two memory tasks. γtACS alone was not administered since a single 3-minute γtACS does not exert any relevant after-effect (Guerra et al., 2018).

*Experiment 2.* We deepen the study of memory effects by confirming experiment 1 results and investigating the course of the memory trace over time after the neuromodulation protocol. We simplified the experimental design by excluding one of the control conditions, i.e., ‘sham-iTBS+sham-γtACS’, as we did not observe any effect related to this condition (see ‘results’ paragraph). In particular, participants were involved in a double-blind cross-over design with two balanced and randomized stimulation conditions (i.e., iTBS+γtACS, iTBS+sham-tACS) separated by a washout week. Participants were required to attend three experimental sessions for each condition. During the first session (day 1), participants received the neuromodulation protocol and then performed FNAT and STMB. In the second session (day 2), participants performed FNAT recall with a 24-hour delay from the neuromodulation protocol, while in the third session (day 7), participants performed FNAT recall and recognition with a 1-week delay (Fig. 1, panel D).

*Experiment 3.* We investigated the neurophysiological effects of the neuromodulation protocol through the combined use of TMS and electroencephalography (TMS-EEG). As for experiment 1, participants underwent an MRI scan to individualize the stimulation sites and permit neuronavigation. Then, they were involved in two balanced, double-blind, randomized experimental sessions of neuromodulation (i.e., iTBS+γtACS, iTBS+sham-tACS) separated by a washout week. TMS-EEG recordings were performed before (T0), immediately after (T1), and 20 minutes after the neuromodulation (T2) (Fig. 1, panel E).

*Experiment 4.* We investigated the effects of the neuromodulation protocol on functional connectivity through fMRI scanning. After the first MRI scanning used for neuronavigation, participants were involved in two balanced, double-blind, randomized experimental sessions of neuromodulation (i.e., iTBS+γtACS, iTBS+sham-tACS) separated by a washout week. The fMRI scanning was performed before (T0) and immediately after (T1) the neuromodulation protocol (Fig. 1, panel F).

### Participants

We recruited young healthy participants who provided written informed consent approved by the Santa Lucia Foundation IRCCS ethical committee (CE/PROG.923) in accordance with the Declaration of Helsinki. See supplementary information for inclusion and exclusion criteria.

In total, 41 subjects were enrolled. 24 participants were involved in experiment 1, in which we tested the effects of the dual iTBS and γtACS on memory functions. Another group of 10 was enrolled in experiment 2 to replicate the results of experiment 1 in an independent sample and to investigate the long-term effect of the dual stimulation. 16 participants were recruited in experiment 3, of which 10 had already participated in experiment 1, to examine the effects on cortical oscillatory activity and cortical reactivity. Finally, 18 participants were involved in experiment 4, of which 12 had already participated in experiment 1, to investigate functional connectivity through fMRI.

Eight participants (i.e., n=4 from experiment 1, n=2 from experiment 3, n=2 from experiment 4) voluntarily withdrew before completing all the experimental sessions. We analyzed at all time points in n=20 subjects for experiment 1, n=10 for experiment 2, n=14 for experiment 3, and n=16 for experiment 4 (see Table 1 for demographic characteristics). See supplementary information for sample size estimation.

**Table 1.** Demographic characteristics.

|  | **Sample**<br>**(n)** | **Age**<br>**(years)\*** | **Sex**<br>**(female\male)** | **Education**<br>**(years)\*** |
| --- | --- | --- | --- | --- |
| **Experiment 1** | 20 | 28.6 (4.0) | 12\8 | 18.8 (3.1) |
| **Experiment 2** | 10 | 24.1 (4.7) | 6\4 | 17.4 (4.7) |
| **Experiment 3** | 14 | 27.1 (2.5) | 11\3 | 19.4 (2.1) |
| **Experiment 4** | 16 | 26.9 (2.4) | 11\5 | 18.9 (1.5) |

\*Mean (s.d.)

### iTBS+_γ_tACS neuromodulation protocol

The co-stimulation consisted of a combination of iTBS and tACS delivered in the gamma band (γtACS) based on previous studies (Guerra et al., 2018; Maiella et al., 2022). The neuromodulation protocol was delivered on the individual PC, based on the individual resting state structural and functional MRI (see ‘MRI data acquisition and preprocessing’ paragraph), and targeted with a stereotaxic neuronavigation system (SofTaxic, E.M.S., Bologna s.r.l.). The active tACS electrode (anode) was placed on the scalp with the iTBS coil above it, and the other tACS electrode (cathode) was placed over the shoulder’s right muscle (Fig. 1, panel C).

tACS was delivered through a Brainstim multifunctional system for low-intensity transcranial electrical stimulation (E.M.S., Bologna s.r.l.) and saline-soaked sponge electrodes (7×5 cm^2^). γtACS sinusoid frequency wave was set at 70 Hz with an intensity of 1mA for a total duration of 190s. iTBS was delivered through a MagStim Rapid^2^ magnetic stimulator (Magstim Company, Whitland, Wales, UK) delivering a biphasic waveform pulse (pulse width _∼_ 0.1 ms) connected to a figure-of-eight coil (70 mm). iTBS consisted of ten bursts of three pulses at 50 Hz lasting 2 s, repeated every 10 s with an 8 s pause between consecutive trains, for a total of 600 pulses lasting 190 s (Huang et al., 2005). Biophysical modeling and E-field calculation were conducted to control for the E-field distribution (see supplementary information). Stimulation parameters were personalized and identified using fMRI, TMS-EEG and electromyography (see supplementary information for details).

No ramp-up and no ramp-down were programmed for the stimulation. Sham stimulation conditions were implemented to control the individual contribution of the techniques and the placebo effect. tACS sham conditions were implemented by applying only a 2s ramp-up and 2s ramp-down at the beginning and the end of stimulation, to give the participant the feeling of real stimulation. Sham-iTBS conditions were implemented by adding a wood layer under the coil(Sandrini et al., 2020). See supplementary information for randomization and masking information.

### Memory tasks

*Face-name associative task.* In experiments 1 and 2, just after the neuromodulation protocol, the participants underwent the FNAT, a cross-modal associative memory test that requires participants to pair pictures of unfamiliar faces with common first names and occupations (Papp et al., 2014). See supplementary information for details about the task construction.

The task consisted of a learning phase followed by a recall and recognition trial. During the learning phase, 12 faces associated with a name and an occupation were shown for 8 seconds on a computer screen. Participants were instructed to read aloud the name and the occupation, to ensure their attention was focused on the items, and to memorize the face-name-occupation association. Right after the learning phase, there was an immediate cued recall phase, in which the participants were asked to recall the name and the occupation of each face previously shown while observing the faces again for 8 seconds each. The delayed cued recall consisted of showing the same faces and asking participants to say each name and occupation. Finally, during the recognition trial, participants were asked to recognize the studied face from a distractor face matched for age and sex. Then, for those names and/or occupations not recollected, they had to identify the correct name and/or occupation among three alternatives each. The distractors were a novel name/occupation and a name/occupation associated with a different face. We considered a correct association when a subject was able to recall all the information for each item (i.e., face, name and occupation), resulting in a total of 36 items to learn and associate. To further investigate the effect on FNAT, we also computed a partial recall score accounting for those items where subjects correctly matched only names with faces (FNAT NAME) and only occupations with faces (FNAT OCCUPATION). See supplementary information for score details. In experiment 1, the learning phase was followed by an immediate cued recall and a 15-minute delayed cued recall with recognition (Fig. S2, panel A in supplementary information) while in experiment 2 the learning phase was followed by an immediate cued recall and a 15-minute delayed recall on day 1, a 24-hour delayed cued recall on day 2, and a 1-week delayed cued recall with recognition on day 7 (Fig. S2, panel B in supplementary information). The recognition trial was administered just in the third session to avoid a learning bias over day 2 and day 7 cued recall.

*The Short-Term Memory Binding test.* This test was run between the FNAT immediate cued recall and the 15-minute delayed cued recall. STMB is a recognition task relying on a change detection paradigm elaborated by Parra and colleagues (Parra et al., 2014). Participants were instructed to remember visual arrays of three black shapes (shape only condition, Fig. S3, panel A in supplementary information) or colored shapes (shape-color binding condition, Fig. S3, panel B in supplementary information) presented for 2 seconds (study phase). After a 1-second delay where a blank screen is presented (retention interval), a display with the same or different items appears in new random locations on the screen (test phase). Participants were asked to press the “1” button on the keyboard if the items shown in the study and test phase were different (50% of the trials) or to press “2” if they were the same. A total of 32 randomized trials were presented for each condition. Conditions were counterbalanced across participants. Before starting the test, each participant underwent a perception trial where the two arrays of shapes were presented on the same screen, to exclude perceptual deficit and to train participants on keyboard answers (Fig. S3, panel C in supplementary information). The test was administered through E-prime 2.0 (Psychology Software Tools, Pittsburgh, USA), which recorded accuracy and reaction times (RTs). We derived two accuracy indexes, respectively for the shape-only condition and the shape-color binding condition (total score of each condition/32x100).

### TMS–EEG data acquisition

Neurophysiological effects in cortical oscillations, excitability, and connectivity were assessed using single-pulse TMS during EEG recordings. During the TMS-EEG assessment, participants sat on a comfortable armchair in a soundproof room, were instructed to fixate on a black cross on the wall, and wore in-ear plugs that continuously played a white noise to avoid possible auditory event-related potential responses (Rocchi et al., 2021). TMS-EEG recordings were performed in a resting state to avoid contamination of cortical activity due to underlying memory tasks. The intensity of the white noise was adjusted individually by increasing the volume (always below 90 decibels) until the participant was sure that s/he could no longer hear the TMS-induced click. TMS for EEG recordings was the same stimulator as for the neuromodulation protocol. The stimulated areas were the PC, the area of interest, and the left posterior parietal cortex (l-PPC), considered as a control area. The order of the stimulation was counterbalanced across participants. PC and l-PPC were both identified by the individual resting state structural and functional MRI (Fig. 1, panel A; see ‘MRI data acquisition and preprocessing’ paragraph). The coil position was constantly monitored using the Softaxic neuronavigation system (E.M.S. Products, Bologna, Italy) and was differently oriented depending on the area of stimulation so that the direction of current flow in the most effective (second) phase was in the posterior-anterior direction. To target the PC, the coil was positioned with an orientation parallel to the midline, while to target the l-PPC, the coil was positioned with an orientation of 15 degrees from the midline (Koch et al., 2018). The single-pulse TMS intensity was set at 100% eSI, separately acquired for individual PC and lPPC (see ‘iTBS+γtACS neuromodulation protocol’ paragraph) without the tACS electrode under the coil. Each TMS-EEG block consisted of 90 single pulses with a randomized inter-stimulus interval (ISI) between 2 and 4 s. EEG was continuously recorded from 61 scalp sites positioned according to the 10-20 International System, using TMS-compatible Ag/AgCl pellet electrodes mounted on an elastic cap (BrainAmp; BrainProducts GmbH, Munich, Germany). Additional electrodes were used as a ground and reference: the ground electrode was positioned in Fpz, while the reference was positioned on the tip of the nose. EEG signals were digitized at a sampling rate of 5 kHz. Skin/electrode impedance was maintained below 5 kΩ. See supplementary information for TMS-EEG preprocessing and analysis.

### MRI data acquisition

MRI data were acquired (1) before experiments 1, 3 and 4, to identify and individualize the stimulation target and (2) before and after the neuromodulation protocol, to assess functional connectivity effects (Fig. 1, Panel C).

Imaging was acquired on a Siemens PRISMA scanner with a 64-channel head-coil (Siemens, USA). During the first acquisition (1), both the structural and functional MRI (fMRI) were run, whereas in the following acquisition (2), only the fMRI was acquired.

The structural imaging was acquired using high-resolution T1-weighted (T1w) anatomical images obtained through a 3D-MPRAGE sequence (TR=2500ms, TE=2 ms, TI=1070ms, flip angle (FA)=8°, thickness=1mm, imaging matrix=240 × 240), and a Diffusion Tensor Images (DTI) sequence Diffusion Tensor Images (DTI) sequence (TR=3400 ms, TE=80ms; 121 directions, b-value=1000 s/mm2, thickness=1.79 mm, gap=1.79 mm, flip angle=90), only acquired before the experiment 3 and 4. The fMRI images were acquired using standard echo-planar blood oxygenation level-dependent (BOLD) imaging (TR=800ms, TE=3 ms, flip angle (FA)=52°, thickness=2.4mm, gap=2.4mm). Subjects were instructed not to focus their thoughts on any particular topic, not to cross their arms or legs, and to keep their eyes open.

For the individualization of the stimulation sites (Fig. 1, panel A), the functional region of interest (ROI) representing the PC node of the DMN (for experiments 1, 3 and 4) and the l-PPC node of the fronto-parietal network (FPN) (for experiment 3) was derived from the Harvard-Oxford atlas available in CONN (Whitfield-Gabrieli and Nieto-Castanon, 2012). A seed-to-voxel correlation map was computed for each participant, thus obtaining a map of positively and negatively correlated voxels, respectively representing the DMN and FPN. An investigator expert in MRI checked both resting-state functional connectivity (rs-FC) maps and structural MRI data (i.e., T1- weighted images) to identify individual hotspots based on ad-hoc criteria. In particular, the stimulation sites were defined as the ones closest to the local maxima of the rs-FC cluster identified as DMN-PC and FPN-PPC being on the top of a cortical gyrus and representing the shortest perpendicular path connecting the stimulating TMS coil on the scalp and the cortex. Based on best judgment, the resulting set of coordinates was picked as the individual stimulation site. The individual set of coordinates was then transformed using a non-linear transformation to reconstruct the targets in individual brain spaces. Lastly, to ensure a coherent intrasession and intersession stimulation (Cocchi and Zalesky, 2018; Fitzgerald et al., 2009), the individualized targets were marked in the subject’s anatomical MRI and loaded into our neuronavigation system. Both personalized points were used for the TMS-EEG assessment, whereas only the personalized PC was used for the neuromodulation protocol in experiments 1, 3 and 4. See supplementary information for fMRI and DTI data preprocessing.

### Statistical analysis

Memory performance and local oscillatory changes data were analyzed through SPSS v22 (IBM, Armonk, NY). Before undergoing parametric or non-parametric statistical procedures, the assumption of normality distribution of data residuals was assessed with Shapiro-Wilk test. The assumption of sphericity was tested with Mauchly test, if this test was significant, we used the Huynh-Feldt correction. The level of significance was set at α=0.05.

To assess memory performances in experiment 1, we run repeated-measures ANOVAs with stimulation condition as a within-subject factor (i.e., iTBS+γtACS; iTBS+sham-tACS; sham-iTBS+sham-tACS) for each dependent variable. In detail, a separate ANOVA was conducted for every FNAT measure (i.e., name, occupation, total) on each memory process (i.e., immediate recall, delayed recall, and recognition). Separate ANOVAs were also conducted for STBM accuracy and RTs both for the shape and the binding condition. Post-hoc comparisons were performed with paired t-tests corrected with Bonferroni method.

To confirm the results obtained from experiment 1 and to investigate the long-lasting effect on long-term memory, we performed experiment 2, where we first ran a repeated-measures ANOVA with stimulation condition as a within-subject factor (i.e., iTBS+γtACS, iTBS+sham-tACS) for the immediate recall, as in experiment 1. Then a two-way repeated measure ANOVA was conducted for FNAT delayed recall performance where stimulation condition (i.e., iTBS+γtACS, iTBS+sham-tACS) and time (i.e., day1, day2, day7) were considered as within-subject factors. Following the results obtained in this analysis (see ‘results’ paragraph), we conducted exploratory paired t-tests aimed at investigating the effect of stimulation conditions on each time point.

Since we do not have an a priori effect size for experiments 1 and 2, we performed a sensitivity power analysis to ensure that these experiments were able to detect the minimum effect size with 80% power and alpha level of 0.05.

To assess cortical oscillation changes in experiment 3, we computed repeated-measures ANOVAs with stimulation condition (iTBS+γtACS, iTBS+sham-tACS) and time T1-T0, T2-T0 (ΔT1, ΔT2) as within factors for each frequency band. Also, in this case, paired t-tests were conducted to investigate the effect of stimulation conditions on each time point. A one-tail hypothesis was considered given the precise hypothesis of gamma increase deriving from our previous work (Maiella et al., 2022).

To assess cortical excitability effects, we used multiple paired t-tests. The analysis was run in the four time windows of the TEP waveform (i.e., 10-30 ms, 21-60 ms, 61-120 ms, 121-150 ms after TMS; see paragraph ‘Cortical excitability’ in ‘TMS-EEG data acquisition and preprocessing’), comparing T1 and T2 to T0 in each stimulation condition. To reduce the occurrence of type I errors, we used the Monte Carlo method, which computes the estimates of the significance probabilities from two surrogate distributions constructed by randomly permuting the two original conditions’ data 3000 times. Moreover, p-values were corrected with the false discovery rate (FDR) method considering the number of electrodes and were considered significant when at least 10 successive t-tests reached the significance threshold (pLJ<LJ0.05). The same statistical analysis was performed to assess changes in source activation. The time series extracted and analyzed from the ROIs (see paragraph ‘source activation’ in ‘TMS-EEG data acquisition and preprocessing’) ranged from -50 to 150 ms from the TMS pulse. As for excitability analysis, we avoided the occurrence of type I errors by implementing Monte Carlo permutation and considering a significant result when at least 10 successive t-tests reached the significance threshold (pLJ<LJ0.05).

To test resting-state functional connectivity (rsFC) modulation, we used a general linear model (GLM). The statistical analyses were carried out using the CONN (v.20b) toolbox and Matlab 2018b software (Mathworks, MA, USA) and were performed considering stimulation conditions (i.e., iTBS+γtACS and iTBS+sham tACS) and time points (i.e., pre and post) as factors. In particular, we performed a ROI-to-ROI analysis focusing on 6 specific ROIs, based on our hypothesis: left and right PPC, medial prefrontal cortex (mPFC), PC, left and right hippocampus, chosen based on Harvard-Oxford (cortical and subcortical) atlas (www.fmrib.ox.ac.uk/fsl/)(Desikan et al., 2006). Specifically, rsFC changes were calculated by computing the Pearson correlation coefficient between the average time series extracted from each individual ROI. Multiple comparisons between ROIs were then performed using a two-sided contrast with a cluster level of p<0.05 FDR-corrected. In addition, the significant nodes were considered as seeds for the voxel-wise analysis, specifically performed by comparing the pre and post iTBS+γtACS. Temporal correlations were calculated between these seeds and all other voxels in the brain. Results were computed by applying a cluster-level threshold of p<0.05 FDR- corrected.

The sample size for experiments 3 and 4 was estimated based on a previous study using the same protocol (Maiella et al., 2022). Based on the effect size reported in this work (η^2^=0.291), our power analysis estimated that a sample size of 14 patients would be necessary to obtain the same effect size with 80% power and an alpha level of 0.05.

To assess linear relationships between the white matter integrity and the functional connectivity outcomes, we perform a bivariate correlation between FA values extracted from the MdLF and the functional connectivity modulation induced by the stimulation assessed through the ROI-to-ROI analysis. Correlations were computed with Pearson’s coefficient (two-tailed).

Data will be available upon rational statement request at https://docs.google.com/spreadsheets/d/1lYQHyXFP2EVT1AwCX-rhndQzPk16Ghb_/edit?usp=sharing&ouid=105875083760241337938&rtpof=true&sd=true.

## Author Contributions

Conceptualization IB, GK; data curation IB, LM; formal Analysis IB, LM, EPC, MA; funding acquisition GK, EPC; investigation IB, MM, LM, MF, FC, ES; methodology IB, LM, MM, EPC, GK; Project administration IB, GK; resources IB, LM, GK; supervision: IB, EPC, SB, GK; visualization: IB, LM, GK; Writing – original draft IB, LM, GK; writing – review & editing IB, LM, MM, SB, EPC, GK.

## Acknowledgments

The authors would like to gratefully acknowledge Maria Stefania De Simone and Marta Rodini for their help in task design and the participants involved in the studies. The draft has been revised by Prof. Alessandro D’Ausilio and Luciano Fadiga.

## Funding resources

The study was founded by H2020 EUROPEAN COMMISSION Future and Emerging Technologies (FET) grant agreement No 101017716 (GK) and the Italian Ministry of University and Research under the National Recovery and Resilience Plan [PE00000006 “A multiscale integrated approach to the study of the nervous system in health and disease” MNESYS (GK) and Fondi DM 502/2022 – PNRR MC42 Bando Giovani Ricercatori (EPC)].

