## Supplementary information for "Dual transcranial electromagnetic stimulation of the precuneus boosts human long-term memory"

**This PDF file includes:**

Supplementary Materials and Methods

References

Figs. S1 to S4

Tables S1 to S4

**Other Supplementary Materials for this manuscript include the following:**

Movies S1

**Supplementary Materials and Methods**

*Enrollment criteria*

Participants were recruited from the Rome metropolitan area via advertisements on local and electronic bulletin boards. Inclusion criteria were young age (18-40 years), native or fluent in Italian, and normal or corrected-to-normal vision and hearing. Exclusion criteria were left-handedness, claustrophobia, psychotropic medication, current or past psychiatric or neurological disorders, history of seizure, metal implant in the head, and implanted electronic device. TMS and tACS safety guidelines and medical regulations were fully followed (Antal et al., 2017; Rossi et al., 2021).

*Sample size estimation*

The sample size for the main experiment 1, 3 and 4 was estimated based on a previous study in which we applied dual iTBS+tACS at different frequencies (theta and gamma) over the dorsolateral prefrontal cortex using TMS-EEG as a read-out of oscillatory cortical activity (Maiella et al., 2022). This study reported an effect size of 0.291 based on a two-way repeated-measures ANOVA in which we compared three neuromodulatory protocols (i.e. iTBS+γtACS; iTBS+shamtACS; iTBS+theta-tACS in three different time points). This analysis revealed a specific increase in gamma activity after the iTBS-γtACS neuromodulatory protocol. Based on this effect size, our power analysis estimated that a sample size of 14 patients would be necessary to obtain the same effect size with 80% power and an alpha level of 0.05.

*iTBS intensity*

To select the intensity for the iTBS protocol we first computed the resting motor threshold (RMT), defined as the lowest intensity producing MEPs of >50 μV in at least five out of 10 trials (Rossini et al., 2015). RMT was tested over the relaxed first dorsal interosseous (FDI) hotspot of the primary motor cortex (M1) in the left dominant hemisphere, with the tACS electrode under the coil to guarantee the same scalp-to-coil distance of the neuromodulation protocol. Electromyographic activity was recorded from the contralateral FDI muscle, using two Ag–AgCl surface cup electrodes (9 mm) in a belly-tendon montage. Responses were amplified through a Digitimer D360 amplifier (Digitimer Ltd, Welwyn Garden City, Hertfordshire, UK). Filters were set at 20 Hz and 2 kHz, with a sampling rate of 5 kHz. iTBS and tACS starting were synchronized using a BrainTrigger (E.M.S., Bologna s.r.l.) and SIGNAL Software. Since the coil-to-cortex distance directly influences the magnitude of magnetic stimulation, for each patient we subsequently calculated a distance-adjusted RMT (AdjRMT). AdjRMT = RMT + m × (DsiteX−DM1) where AdjRMT is the adjusted MT in % of stimulator output, MT is the unadjusted MT in % of stimulator output, DM1 is the distance between the scalp and M1 hotspot, DSiteX is the distance between the scalp and a second cortical region (SiteX), and m is the distance-effect gradient (Stokes et al., 2007). Afterward, to optimize the stimulation intensity, each patient received 50 TMS single pulses at an initial intensity of 100% of adjMT over the individualized PC during a 64-channel EEG recording, which permitted the visualization of TMS-evoked potential (TEPs)(Mancuso et al., 2021). The intensity of TMS was eventually increased in steps of 2% of the maximal stimulator output based on the visualization of a first TEP peak of at least 3 µV (Koch et al., 2022) reaching the effective stimulation intensity of PC (PC-eSI). Finally, the stimulation intensity for iTBS was set at 80% of the PC-eSI.

*Randomization and masking*

Memory tasks, TMS-EEG and fMRI were performed by researchers blinded to neuromodulation condition (memory task: I.B., F.C. and E.S.; neurophysiological data: I.B. and M.M.; fMRI: L.M.). The neuromodulation protocols were performed by a dedicated technician (M.F). All subjects received every stimulation condition thanks to the cross-over design. The stimulation conditions orders were randomized and balanced among subjects. Moreover, in experiments 1 and 2, the parallel forms of the memory task were equally distributed among the stimulation conditions orders.

*Biophysical modeling and E-field calculation*

We calculated the norm E-field distribution induced by tACS and iTBS over the PC for each participant included in experiments 1, 3 and 4 using the simulation package SimNIBS (version 4.0) (Fig. 1, panel B). The realistic head model was built from the structural T1 and T2-weighted MRI of each single-subject. Default isotropic conductivities were used in our simulation (Thielscher et al., 2015). The final mesh was comprehensive of grey (GM) and white matter (WM), scalp tissue, bone, and cerebrospinal fluid (CSF) (see (Windhoff et al., 2013) for further modeling details). Each tissue segmentation was carefully examined slice-by-slice to ensure proper classification.

The TMS E-field distribution was computed using the model of the Magstim 70 mm figure-of-eight coil, used for the iTBS (Thielscher and Kammer, 2002). We calculated the E-field input in the form of dI/dt in units of A/us based on the coil model, stimulator model, and pulse intensity for each subject (for details see (Kammer et al., 2001)). The center of the coil was positioned over the target area, expressed in individual MNI coordinates, as extracted from baseline MRI. The coil model was then manually moved and rotated according to the specified position and direction used in the real stimulation. The tACS E-field distribution was calculated using the 7x5 rectangular sponge electrodes with an electrode thickness of 2 mm and a sponge thickness of 4 mm, centered in the target area of stimulation. Moreover, using an ad-hoc script we simulated the E-field distribution of the combined techniques (iTBS+tACS) for each subject.

The output is reported in Table S4 and is represented in V/m. Figure S1 also shows TMS and tACS E-field calculation for a representative subject.

*Face-name associative task construction and scoring*

For this study, FNAT was adapted from an Italian version to create three parallel forms by selecting images from an online dataset (i.e., FACES, Center for Lifespan Psychology, Max Planck Institute for Human Development, Berlin, Germany) and pairing them with Italian names and occupations (De Simone et al., 2022; Ebner et al., 2010). Each picture's face size was 19x15cm. In the learning phase, faces were shown along with names and occupations for 8 seconds each (totaling approximately 2 minutes). During immediate recall, the faces were displayed alone for 8 seconds. In the delayed recall and recognition phase, pictures were presented until the subject provided answers. We used a different set of stimuli for each stimulation condition, resulting in a total of 3 parallel task forms balanced across conditions and session order. All parallel forms comprised 6 male and 6 female faces; for each sex, there were 2 young adults (around 30 years old), 2 middle-aged adults (around 50 years old), and 2 elderly adults (around 70 years old). Before the experiments, we conducted a pilot study to ensure no differences existed between the parallel forms of the task. The task and its parallel form can be provided upon request.

The score was computed by deriving an accuracy percentage index dividing by 12 and multiplying by 100 the correct association sum. The partial recall scores were computed in the same way only considering the sum of face-name (NAME) and face-occupation (OCCUPATION) correctly recollected.

Each accuracy percentage index was computed for each task phase: immediate cued recall, delayed cued recall, and recognition for experiment 1; immediate cued recall, delayed cued recall (day 1), 24-hours delayed cued recall (day2), 1-week cued recall and recognition (day7) for experiment 2.

*TMS–EEG preprocessing and analysis*

*EEG preprocessing.* TMS-EEG data were preprocessed offline with Brain Vision Analyzer (Brain Products GmbH, Munich, Germany). Data were segmented into epochs starting 1 s before and 1 s after the TMS pulse. TMS pulse artifact was removed and replaced using cubic interpolation, from 1 ms before to 10 ms following the pulse. Afterward, data were down-sampled to 1,000 Hz and band-pass filtered between 1 and 80 Hz (Butterworth zero-phase filters). A 50-Hz notch filter was applied to reduce noise from electrical sources. Then, all the epochs were visually inspected and those with an excessively noisy EEG were excluded from the analysis. Independent component analysis (INFOMAX-ICA) was applied to the EEG signal to identify and remove components reflecting muscle activity, eye movements, blink-related activity, and residual TMS-related artifacts based on previously established criteria (Casula et al., 2017; Hernandez-Pavon et al., 2023). Finally, the signal was re-referenced to the average signal of all the electrodes.

*Source activation.* To characterize cortical source activation at the individual anatomical level, each participant’s MRI was uploaded on Brainstorm toolbox, segmented through CAT12 (Christian Gaser et al., 2022), and normalized. TMS-EEG recordings were then uploaded for each subject and condition, adjusting the channel montage over the cortical surface extract from the individual MRI. The head model was computed using the symmetric boundary element method (symmetric BEM) running on Open MEEG (Gramfort et al., 2010; Kybic et al., 2005). Noise covariance was calculated referring to a baseline ranging from -500 to -1 ms relative to the TMS pulse and removing DC offset block by block. Source activity was computed adopting the minimum norm imaging method, the current density map was set as measure and a constrained dipole orientation was selected. The estimated sources were transformed in z-score points considering a baseline window of -100 to -1 ms and were then spatially smoothed (SurfStat, KJ, Worsley) with a full width at half maximum value set at 3 mm. Once source activity was calculated, we projected it to the default ICBM152 cortex surface to permit group visualization (Movie S1).

*Cortical oscillations.* The analysis of cortical oscillations was performed running a time/frequency decomposition based on Morlet wavelet (cycles: 5; frequency resolution: 1 Hz from 4 to 90 Hz; baseline correction -700;-300 ms) and computing the TMS-related spectral perturbation (TRSP) for theta, alpha, beta, and gamma band (Casula et al., 2022, 2018). To analyze local oscillatory activity, we computed the average TRSP within the following clusters of electrodes: Pz, P1, P2, POz for PC; P3, P5, CP3, PO3 for l-PPC. For each time, condition, and stimulated area, we computed the corresponding cluster, and averaged the TRSP values between 10 and 250 ms (i.e., the average time window of the spectral perturbation) in the following frequency bands: theta (4-7 Hz), alpha (8-13 Hz), beta (14-30 Hz), low-gamma (31-40 Hz), high-gamma (41-90 Hz).

*Cortical excitability.* Changes in cortical excitability were analyzed through the TMS-evoked potentials (TEPs) by importing the pre-processed EEG data in Brainstorm toolbox (Tadel et al., 2011) running in a MATLAB environment (MathWorks Inc., Natick, MA). TEPs were computed by averaging all the time-locked EEG responses in each electrode from 100 ms before to 150 ms after the TMS pulse, with a baseline correction of 100 ms before the TMS pulse. TEPs amplitude was then computed as the mean activity within four time windows (W) based on an accurate visual inspection (Casula et al., 2023; Koch et al., 2018): W1 from 10 to 30 ms, W2 from 31 to 60 ms, W3 from 61 to 120 ms, and W4 from 121 to 145 ms after the TMS pulse.

*fMRI data preprocessing.* SPM12 (Statistical Parametric Mapping), CONN, and MATLAB 2018a (MathWorks, MA, USA) software were used to pre-process datasets. The following preprocessing steps were applied to the BOLD images: discarding of the first three volumes to allow for steady-state magnetization and stabilization of participant status; slice timing; realigning to correct for head motion; co-registration to structural images; segmentation; nonlinear normalization to the Montreal Neurological Institute (MNI) template brain; voxel resampling to an isotropic 3 × 3 × 3 mm voxel size; smoothing with an isotropic Gaussian kernel (full-width at half maximum, 8 mm). Structural images were co‐registered to the mean volume of functional images and segmented. Linear trends were removed to reduce the influence of the rising temperature of the MRI scanner and all functional volumes were bandpass‐filtered at 0.01 Hz < f < 0.08 Hz to reduce low‐frequency drifts. Finally, we regress out potential confounding signals, like physiological high-frequency respiratory and cardiac noise, from grey matter voxels’ BOLD time course using the Compcorr algorithm (Whitfield-Gabrieli and Nieto-Castanon, 2012), to reduce artificial negative correlation and provide adequate filtering of the data.

*DTI preprocessing*. The FMRIB Software Library (FSL) and DSI-studio were used to pre-process datasets. Binary masks were made and then the data were corrected for susceptibility artifacts using the TOPUP function. Volumes acquired with opposing phase-encoding directions were combined and then corrected for artifacts of motion and eddy-current distortion using EDDY. The corrected diffusion-weighted dataset and reoriented b-table were then imported to DSI Studio, registered (rigid body), and resampled to the space of T1. The accuracy of b-table orientation was examined by comparing fiber orientations with those of a population-averaged template (Yeh et al., 2018). The restricted diffusion was quantified using restricted diffusion imaging (Yeh et al., 2017). The diffusion data were reconstructed using generalized q-sampling imaging (GQI; (Yeh et al., 2010) with a diffusion sampling length ratio of 1.25. The tensor metrics were calculated using DWI with a b-value lower than 1750 s/mm². A deterministic fiber tracking algorithm (Yeh et al., 2013) was used with augmented tracking strategies (Yeh, 2020) to improve reproducibility. The anatomy prior of a tractography atlas (Yeh et al., 2018) was used to map the bilateral Middle Longitudinal Fasciculus (MdLF) with a distance tolerance of 16 mm in the ICBM152 space. The anisotropy threshold was randomly selected. The angular threshold was randomly selected from 15 degrees to 90 degrees. The step size was randomly selected from 0.5 to 1.5 voxels. Tracks with lengths shorter than 30 or longer than 300 mm were discarded. A total of 10000 tracts was calculated. Topology-informed pruning (Yeh et al., 2019) was applied to the tractography with 16 iteration(s) to remove false connections. Finally, the MdLF fractional anisotropy (FA) was extracted for each subject.

**Supplementary Figures**

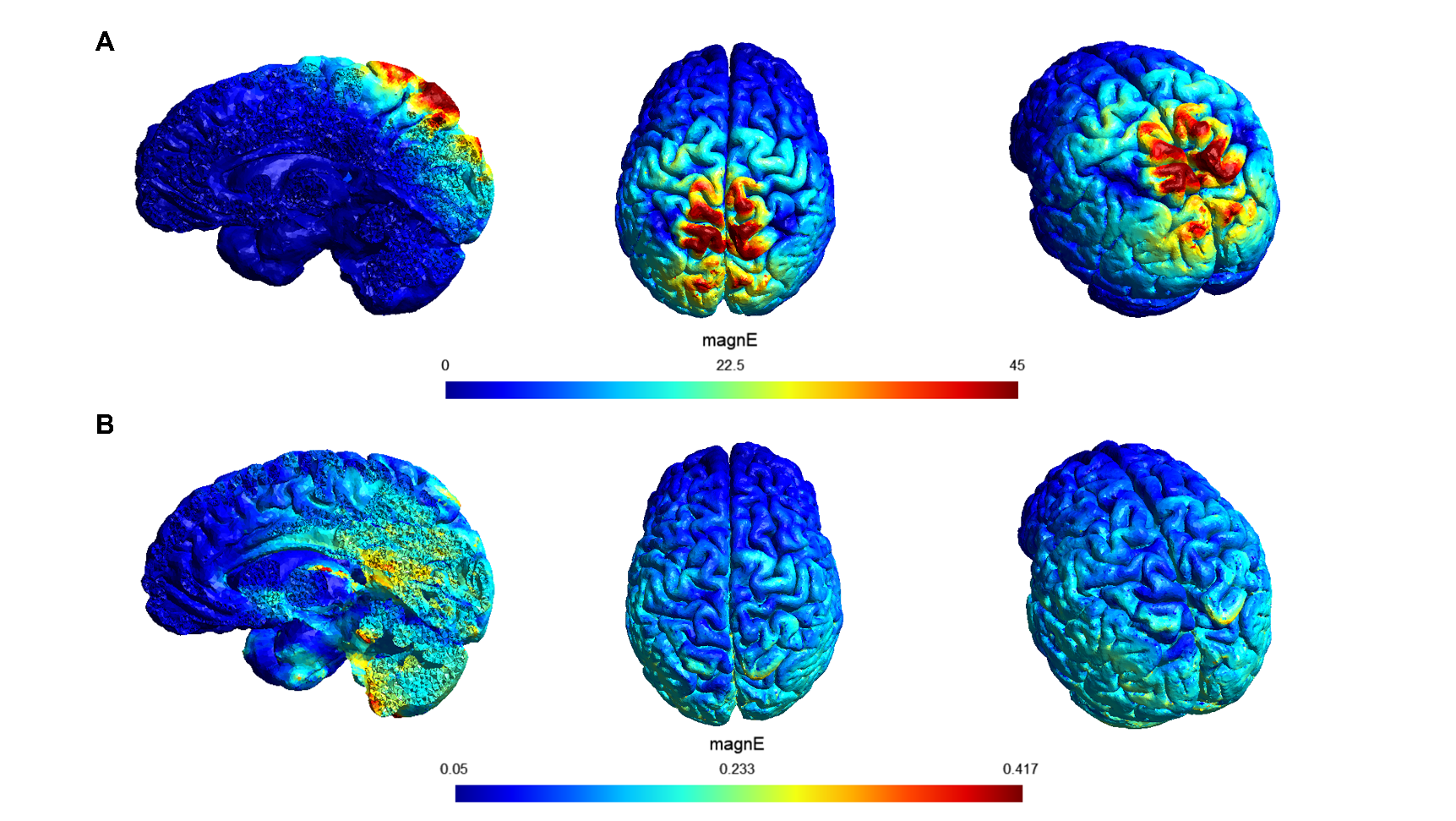

Fig. S1. TMS and tACS e-field simulation.

Figure S1 shows the e-field for a representative subject. Panel (**A**) shows TMS e-fiel, while panel (**B**) shows tACS e-field simulation.

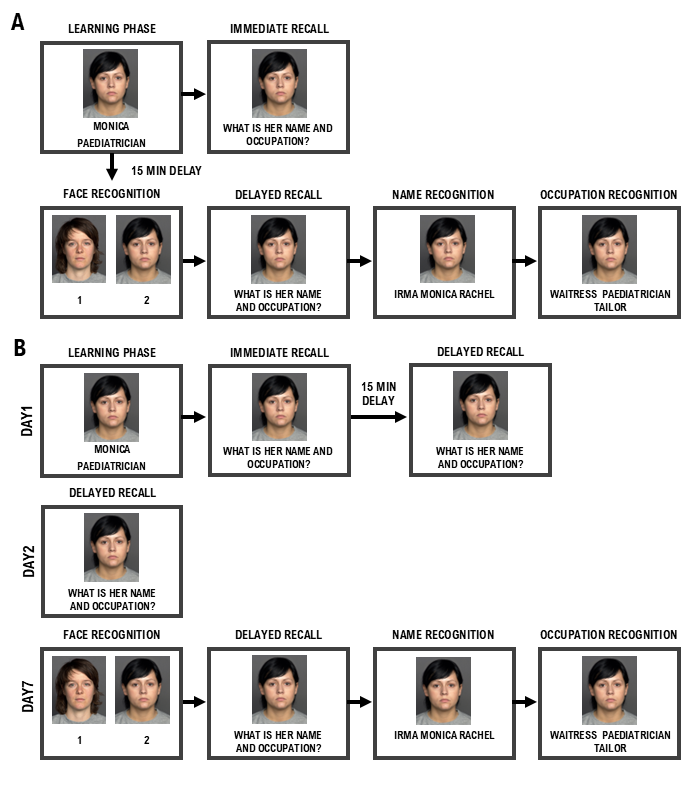

Fig. S2. Face-name associative task (FNAT).

Panel (**A**) shows FNAT steps implemented in experiment 1consisting of a learning phase followed by immediate cued recall and a 15-minute delayed cued recall with recognition. Panel (**B**) shows the variation of FNAT implemented in experiment 2, where the learning phase was followed by an immediate cued recall and a 15-minute delayed recall on day 1, a 24-hour delayed cued recall on day 2, and a 1-week delayed cued recall with recognition on day 7.

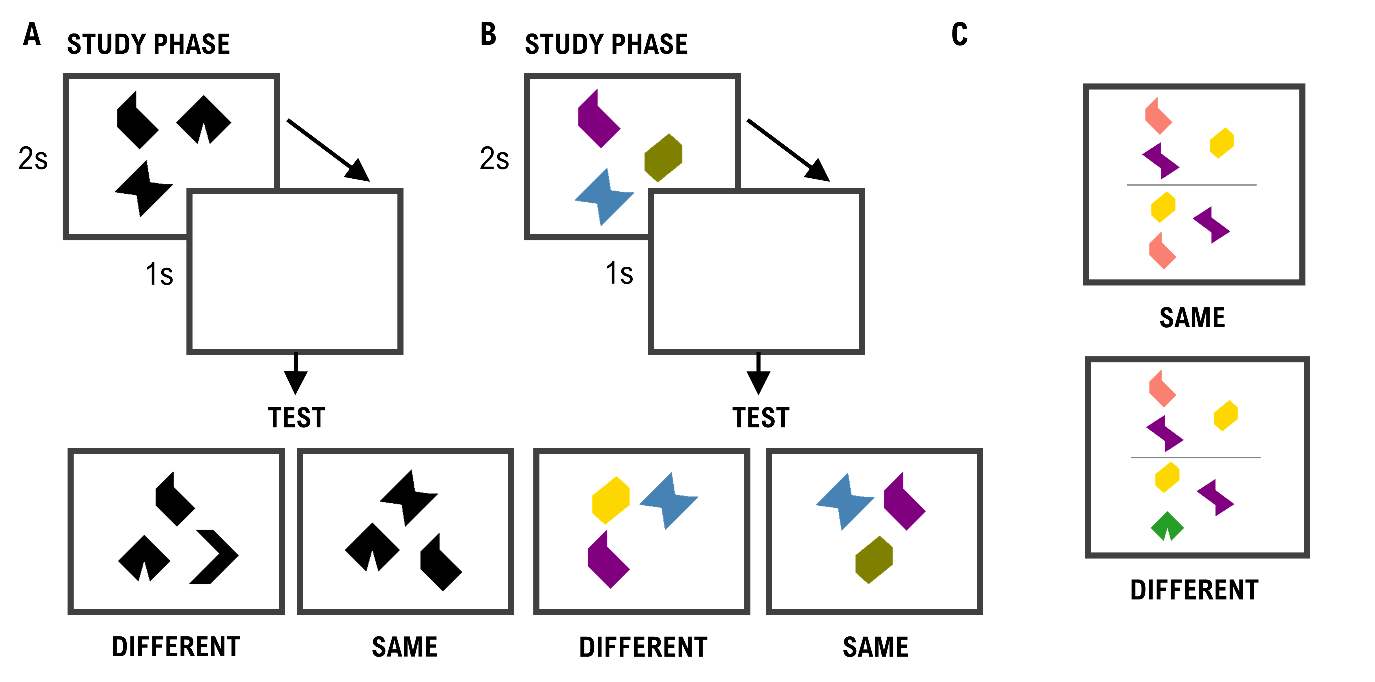

Fig. S3. Short-term memory binding task (STMB).

Panel (**A**) represents the STMB for the shapes only, while panel (B) represents STMB for shapes and colors. Both versions consist of a study phase (up) lasting 2s, followed by a retention phase (center) lasting 1s, and a test phase (down) where the participants have to decide if the items shown are the same or different to the learning phase. Panel (**C**) shows the perception trial, where two arrays of shapes are presented at the same time and participants have to answer if they are the same or different.

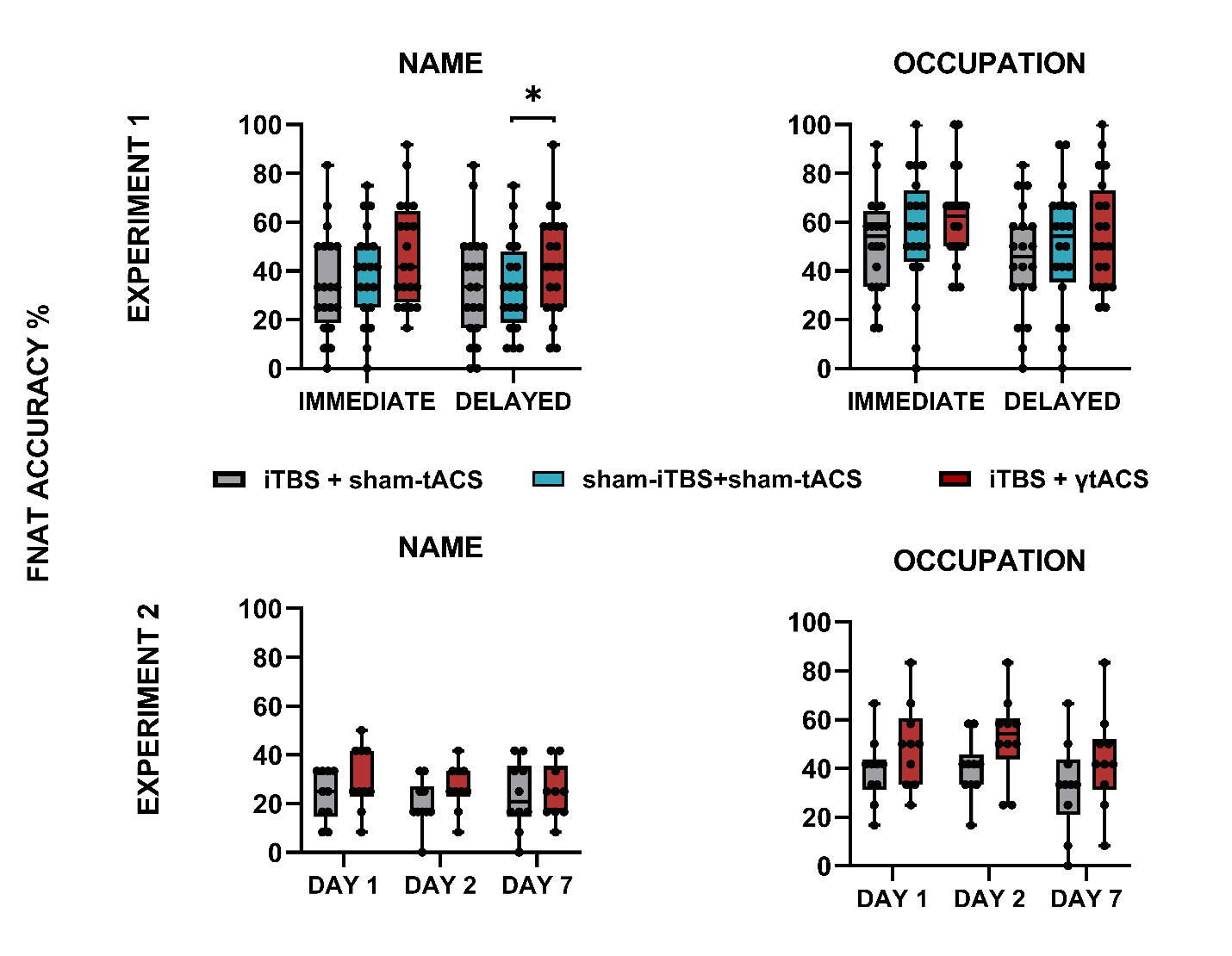

**Fig. S4. FNAT in-depth analysis.** In-dept Face-name associative task (FNAT) accuracy for the association of face with name (NAME) and face with occupation (OCCUPATION). The first raw shows the performances in immediate and delayed trials resulting from experiment 1 (N=20). The second raw shows the performances in the delayed trials resulting from experiment 2 (N=10). Boxes delimit the lower (Q1) and the upper (Q3) quartile, whiskers extend to the smallest and largest values, dots represent individual values. Grey corresponds to iTBS+sham-tACS, blue to sham-iTBS+sham-tACS and red to iTBS+γtACS. * = p<0.05

Table S1.

Experiment 1 statistical details of FNAT and STMB test.

| **Outcome measure** | **Mean (sd)** | | | **Stimulation effect** | |
| --- | --- | --- | --- | --- | --- |
|  | **iTBS+sham-tACS** | **sham-iTBS+sham-tACS** | **iTBS+γtACS** | **F_df_** | **p** |
| **FNAT immediate** | | | | | |
| TOTAL | 25 (17.9) % [3.0 (2.2)] | 30.8 (21.1) % [3.7 (2.5)] | 40 (21.6) % [4.8 (2.6)] | 7.190 _2,38_ | 0,002 |
| NAME | 34.6 (21.2) % [4.2 (2.5)] | 38.8 (19.9) % [4.7 (2.4)] | 46.7 (21.2) % [5.6 (2.5)] | 3.200 _2,38_ | 0,052 |
| OCCUPATION | 50.8 (20.2) % [6.1 (2.4)] | 55.8 (24.9) % [6.7 (3.0)] | 61.3 (19.7) % [7.4 (2.4)] | 2.610 _2, 38_ | 0,086 |
| **FNAT delayed** | | | | | |
| TOTAL | 24.2 (19.8) % [2.9 (2.4)] | 26.3 (17.6) % [3.2 (2.1)] | 34.2 (20.4) % [4.1 (2.4)] | 5.860 _2,38_ | 0,006 |
| NAME | 33.8 (22.7) % [4.1 (2.7)] | 33.8 (19) % [4.1 (2.3)] | 42.9 (21.5) % [5.2 (2.46] | 3.460 _2,38_ | 0,042 |
| OCCUPATION | 44.2 (22) % [5.3 (2.6)] | 50 (25.5) % [6.0 (3.1)] | 55 (22.7) % [6.6 (2.7)] | 2.880 _2,38_ | 0,068 |
| **FNAT recognition** | | | | | |
| TOTAL | 61.7 (24.4) % [7.4 (2.9)] | 65 (20) % [7.8 (2.4)] | 65 (16.6) % [7.8 (2.0)] | 0.401 _2,38_ | 0,673 |
| NAME | 66.7 (23.4) % [8.0 (2.8)] | 70.8 (16.8) % [8.5 (2.9)] | 71.7 (14.4) % [8.6 (1.7)] | 0.694 _2,38_ | 0,506 |
| OCCUPATION | 84.6 (15.4) % [10.2 (1.8)] | 88.3 (13.6) % [10.6 (1.6)] | 84.6 (10.9) % [10.2 (1.3)] | 0.733 _2,38_ | 0,487 |
| **STMBT RT** | | | | | |
| shape | 1550 (341) ms | 1571 (513) ms | 1597 (503) ms | 0.188 _2,38_ | 0,829 |
| binding | 1652 (363) ms | 1731 (446) ms | 1809 (454) ms | 2.660 _2,38_ | 0,083 |
| **STMBT accuracy** | | | | | |
| shape | 96.5 (4.2) % | 95.8 (4.67) % | 96 (3.12) % | 0.172 _2,38_ | 0,842 |
| binding | 78 (10.2) % | 78.8 (11.1) % | 77.8 (9.21) % | 0.062 _2,38_ | 0,94 |

[ ] represent raw scores.
Table S2.

Experiment 2 statistical details of immediate and recognition FNAT and STMB test.

| **Outcome measure** | **mean (sd)** | | **Stimulation effect** | |
| --- | --- | --- | --- | --- |
|  | **iTBS+shamtACS** | **iTBS+γtACS** | **F_df_** | **p** |
| **FNAT immediate** | 17.5 (8.29) % [2.1 (1.0)] | 26.7 (10.2) % [3.2 (1.2)] | 7.310 _1,9_ | 0,024 |
| **FNAT recognition** | 43.3 (15.6) % [5.2 (1.9)] | 46.7 (21.9) % [5.6 (2.6)] | 0.255 _1,9_ | 0,625 |
| **STMBT RT** | | | | |
| shape | 1493 (324) | 1547 (389) | 0.348 _1,9_ | 0,570 |
| binding | 1906 (352) | 1803 (538) | 1.100 _1,9_ | 0,323 |
| **STMBT accuracy** | | | | |
| shape | 95.5 (3.81) | 96.1 (4.25) | 0.310 _1,9_ | 0,591 |
| binding | 70.6 (12.5) | 76.5 (9.71) | 2.700 _1,9_ | 0,135 |

[ ] represent raw scores.

Table S3.

Experiment 2 statistical details of the delayed FNAT.

|  | **DAY1 mean (sd)** | | **DAY2 mean (sd)** | | **DAY7 mean (sd)** | | **Stimulation effect** | | **Delay effect** | | **Stimulation * Delay effect** | |
| --- | --- | --- | --- | --- | --- | --- | --- | --- | --- | --- | --- | --- |
| **Outcome measure** | **iTBS+sham**  **tACS** | **iTBS+γtACS** | **iTBS+sham**  **tACS** | **iTBS+γtACS** | **iTBS+sham**  **tACS** | **iTBS+γtACS** | **F_df_** | **p** | **F_df_** | **p** | **F_df_** | **p** |
| **FNAT delayed** | 15 (9.46) % [1.8 (1.1)] | 25.8 (10.7) % [3.1 (1.3)] | 14.2 (9.66) % [1.7 (1.2)] | 25 (9.62) % [3.0 (1.2)] | 14.2 (12.5) % [1.7 (1.5)] | 23.3 (12.9) % [2.8 (1.5)] | 8.433 _1,9_ | 0,017 | 0.419 _2,18_ | 0,664 | 0.286 _2,18_ | 7,55 |

[ ] represent raw score

Table S4.

Subject-specific information about stimulation parameters and e-field calculations.

| **Code** | **Experiment (n)** | **75% eSI (%)** | **Coil-to-cortex distance PC (mm)** | **PC individualized coordinate (x,y,z)** | | | **TMS e-field (V/m)** | **tACS e-field (V/m)** | **TMS+tACS e-field (V/m)** |
| --- | --- | --- | --- | --- | --- | --- | --- | --- | --- |
| 001C | 1,3,4 | 42 | 17.59 | 9 | -31 | 40 | 19 | 0.1 | 19 |
| 002P | 1,3,4 | 46 | 19.75 | -1 | -20 | 56 | 21 | 0.09 | 21 |
| 003M | 1 | 51 | 24.23 | -3 | -67 | 51 | 27 | 0.17 | 25 |
| 004S | 1,3,4 | 44 | 18.62 | 5 | -40 | 50 | 33 | 0.11 | 33 |
| 006L | 1,4 | 43 | 14.61 | 5 | -36 | 56 | 33 | 0.13 | 33 |
| 007S | 1 | 44 | 19.19 | 5 | -32 | 45 | 31 | 0.11 | 31 |
| 008F | 3,4 | 46 | 19.65 | 2 | -47 | 30 | 37 | 0.12 | 36 |
| 009D | 1 | 43 | 18.19 | -6 | -74 | 44 | 25 | 0.2 | 25 |
| 010P | 1 | 39 | 20.09 | -2 | -68 | 52 | 21 | 0.19 | 21 |
| 011V | 1,3,4 | 43 | 20.74 | -1 | -22 | 48 | 30 | 0.11 | 30 |
| 012M | 1 | 44 | 19.82 | 1 | -58 | 63 | 28 | 0.11 | 29 |
| 013D | 1 | 37 | 24.19 | 5 | -75 | 61 | 34 | 0.16 | 34 |
| 014C | 1 | 45 | 19.24 | -3 | -63 | 56 | 32 | 0.13 | 32 |
| 015G | 1 | 44 | 19.53 | 4 | -63 | 57 | 34 | 0.24 | 34 |
| 016L | 1 | 44 | 16.89 | 1 | -66 | 50 | 35 | 0.22 | 35 |
| 017M | 1 | 41 | 19.94 | 10 | -78 | 46 | 28 | 0.14 | 28 |
| 018B | 3,4 | 56 | 24.16 | 0 | -31 | 68 | 22 | 0.11 | 23 |
| 019B | 1 | 45 | 18.99 | 5 | -75 | 41 | 26 | 0.18 | 26 |
| 020D | 1,3,4 | 45 | 16.16 | 7 | -60 | 10 | 38 | 0.24 | 38 |
| 022B | 1,3 | 55 | 20.11 | 2 | -52 | 34 | 31 | 0.17 | 31 |
| 023R | 1,3,4 | 48 | 23.23 | 0 | -49 | 31 | 30 | 0.14 | 29 |
| 024A | 1,3 | 45 | 20.12 | 7 | -41 | 45 | 35 | 0.11 | 35 |
| 035R | 4 | 53 | 18 | -2 | -39 | 39 | 26 | 0.16 | 26 |
| 036M | 4 | 65 | 24.99 | -2 | -30 | 47 | 35 | 0.07 | 35 |
| 037I | 3,4 | 37 | 17.04 | 3 | -37 | 58 | 30 | 0.14 | 30 |
| 038P | 3,4 | 42 | 19.09 | 4 | -31 | 26 | 14 | 0.16 | 14 |
| 039C | 3,4 | 49 | 16.5 | -2 | -36 | 23 | 22 | 0.14 | 22 |
| 040A | 3,4 | 54 | 20.27 | 2 | -50 | 46 | 48 | 0.09 | 48 |
| 041A | 4 | 52 | 21.54 | 0 | -29 | 34 | 25 | 0.09 | 25 |

Movie S1.

The movie shows the source reconstruction of the TMS-EEG signal when targeting the precuneus. The three columns correspond to the recordings conducted before, just after and after 20 minutes from the non-invasive brain stimulation protocol (i.e., T0, T1, and T2). The first row corresponds to the iTBS+γtACS condition while the second to the iTBS+shamtACS condition.
