## Supplementary figures and images for "Dual transcranial electromagnetic stimulation of the precuneus boosts human long-term memory"

### figure 1

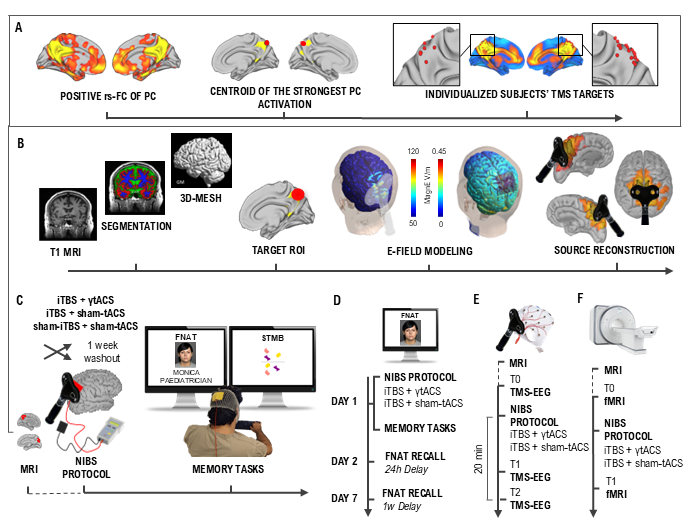

### figure 3

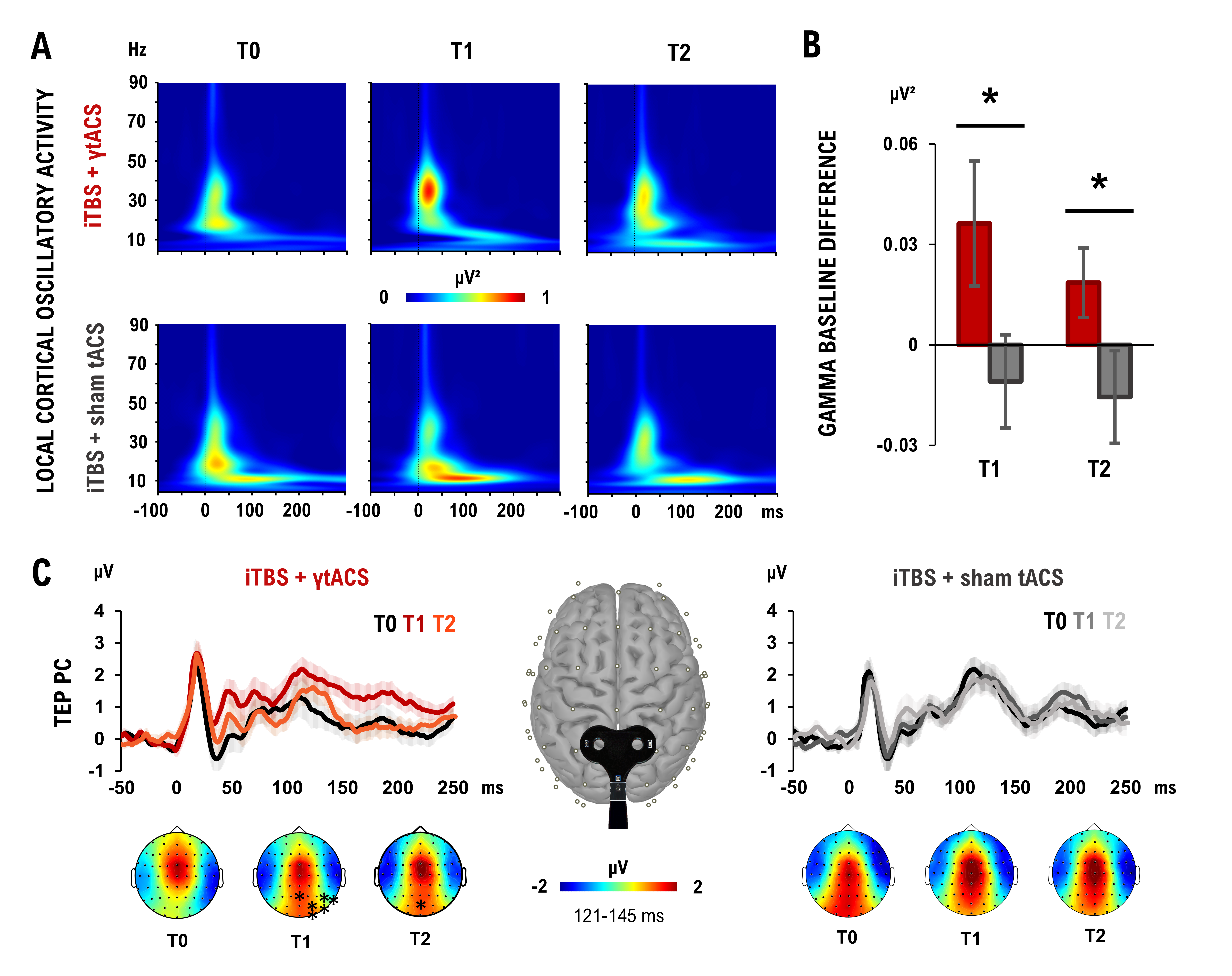

### figure 4

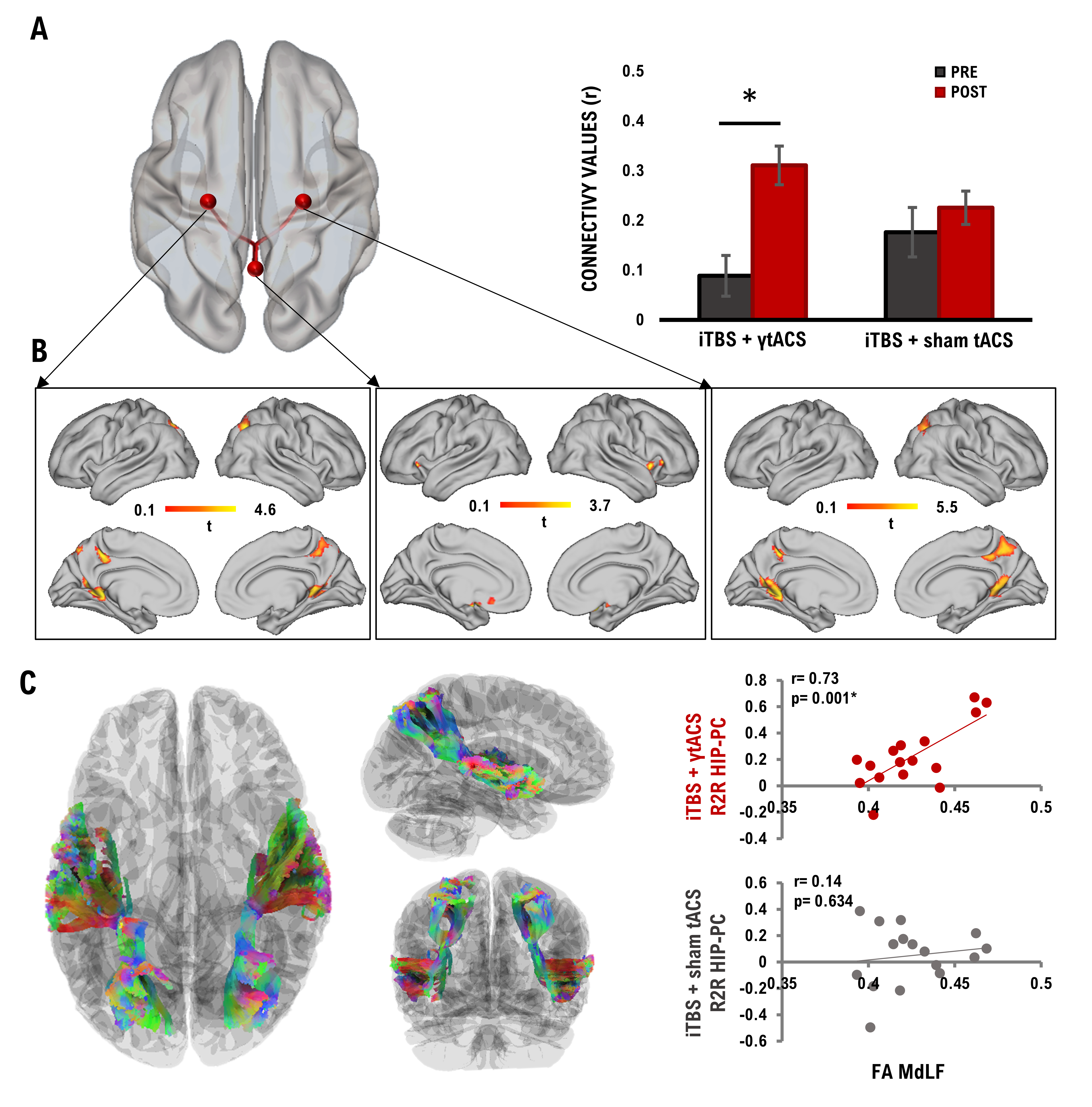

### figure S1

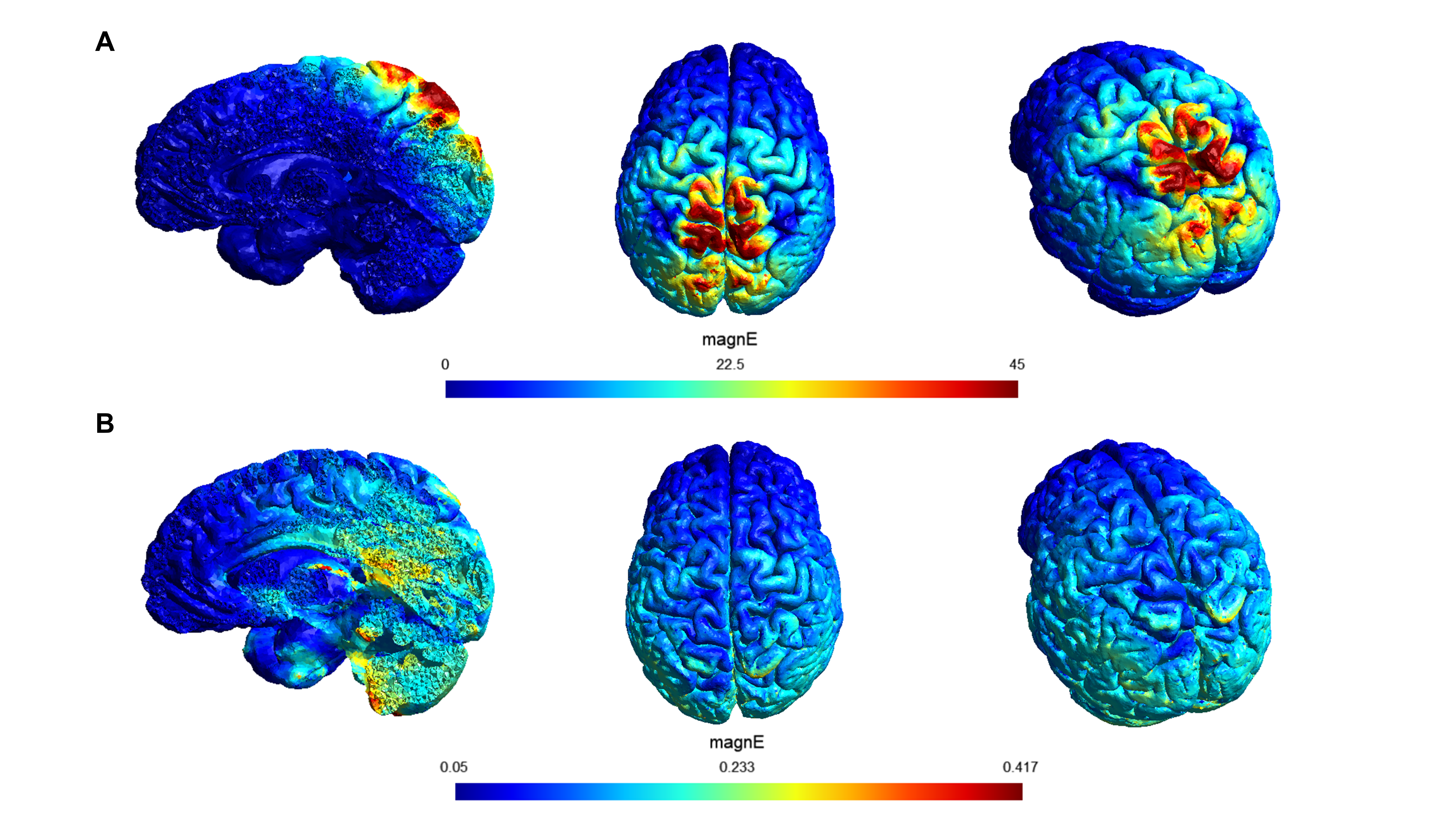

### figure S2

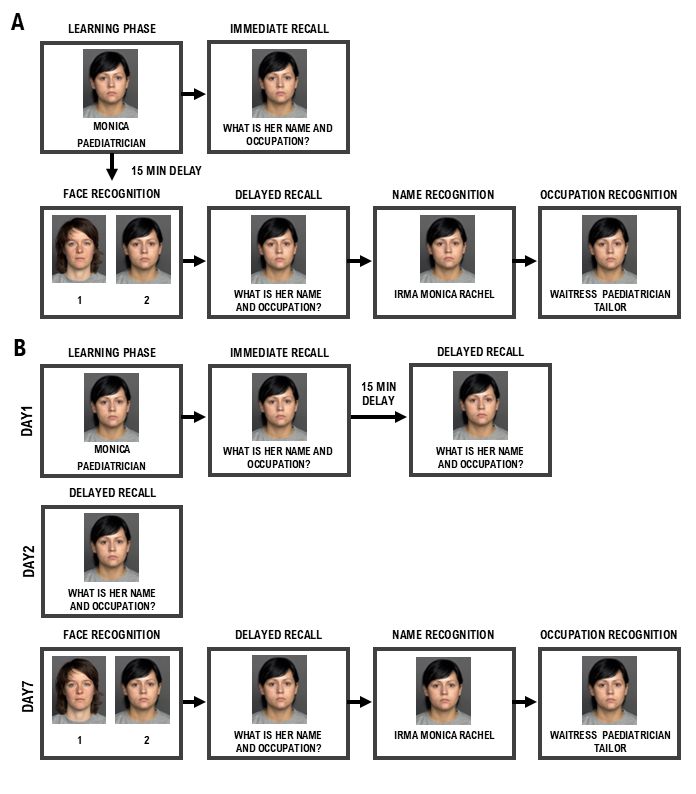

### figure S3

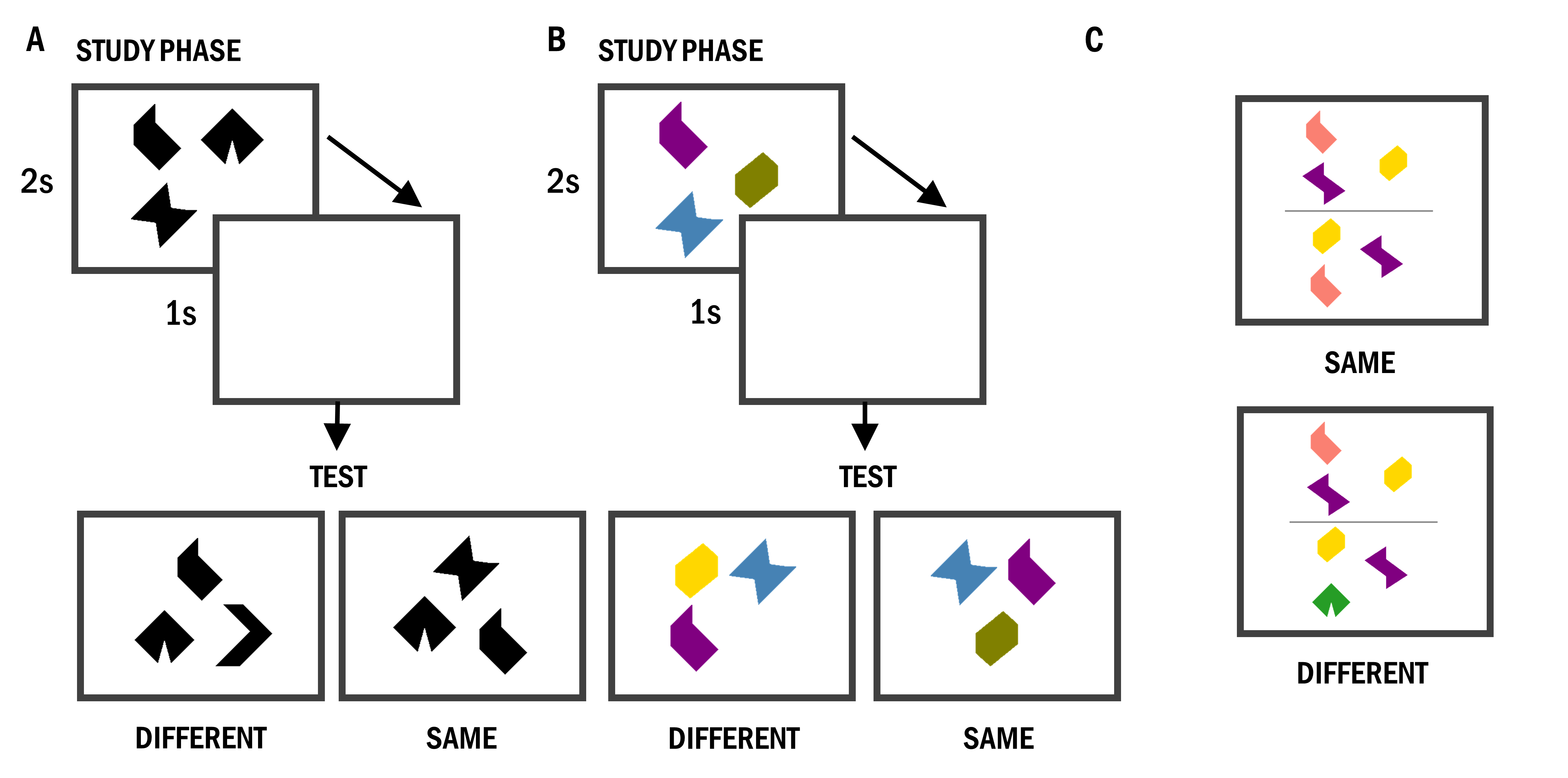

### figure S4

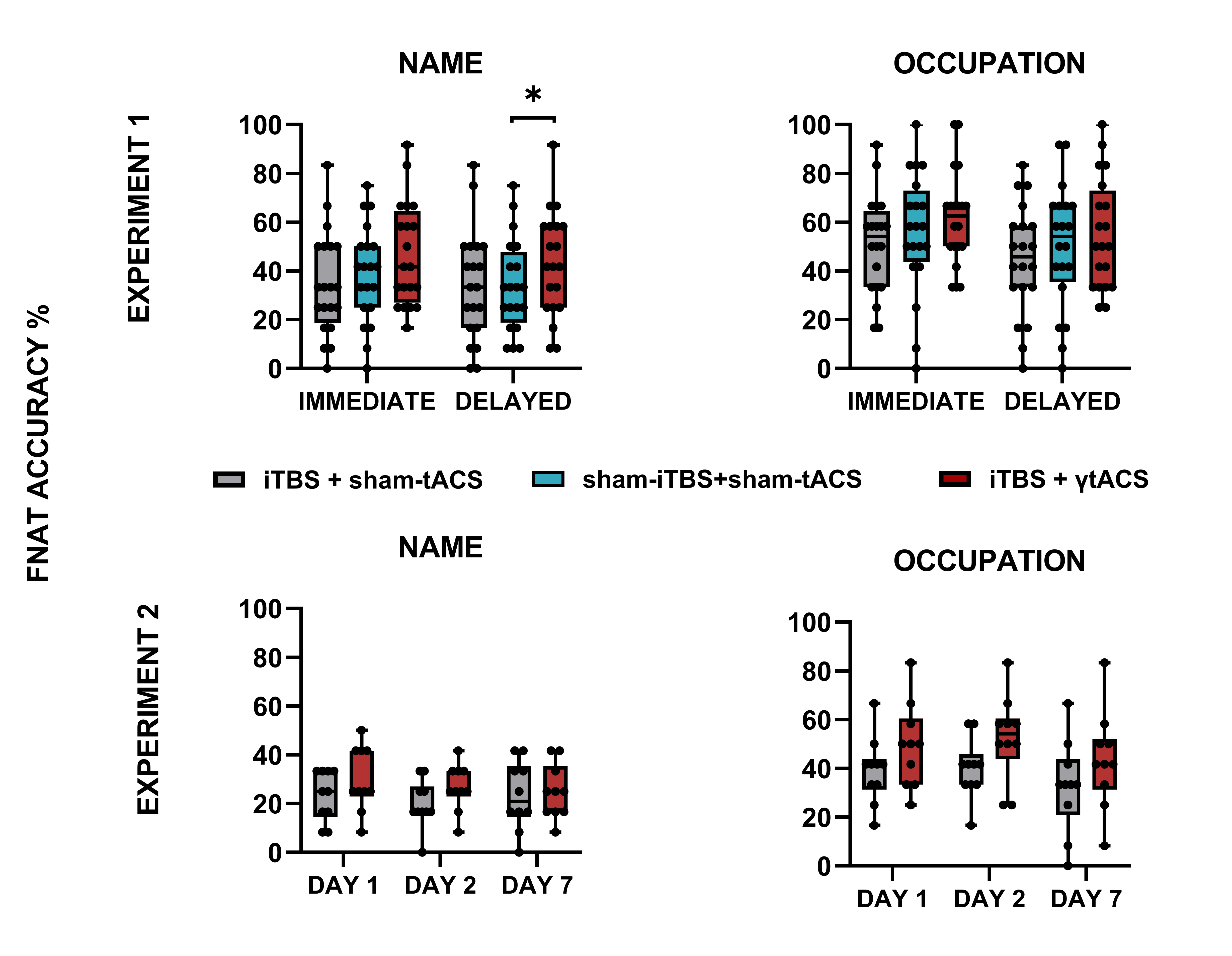
